# Rapid auxin-mediated phosphorylation of Myosin regulates trafficking and polarity in Arabidopsis

**DOI:** 10.1101/2021.04.13.439603

**Authors:** Huibin Han, Inge Verstraeten, Mark Roosjen, Ewa Mazur, Nikola Rýdza, Jakub Hajný, Krisztina Ötvös, Dolf Weijers, Jiří Friml

## Abstract

The signaling molecule auxin controls plant development through a well-known transcriptional mechanism that regulates many genes. However, auxin also triggers cellular responses within seconds or minutes, and mechanisms mediating such fast responses have remained elusive. Here, we identified an ultrafast auxin-mediated protein phosphorylation response in Arabidopsis roots that is largely independent of the canonical TIR1/AFB receptors. Among targets of this novel response are Myosin XI and its adaptor protein MadB2. We show that their auxin-mediated phosphorylation regulates trafficking and polar, subcellular distribution of PIN auxin transporters. This phosphorylation-based auxin signaling module is indispensable during developmental processes that rely on auxin-mediated PIN repolarization, such as termination of shoot gravitropic bending or vasculature formation and regeneration. Hence, we identified a fast, non-canonical auxin response targeting multiple cellular processes and revealed auxin-triggered phosphorylation of a myosin complex as the mechanism for feedback regulation of directional auxin transport, a central component of auxin canalization, which underlies self-organizing plant development.

## Introduction

The plant hormone auxin acts as a key regulator of plant development and controls multiple cellular processes. The current main paradigm of auxin action is that it transcriptionally reprograms individual cell behavior to continually perpetuate and adapt overall plant development (Bargman et al., 2013; Weijers and Wagner, 2016; Lavy and Estelle, 2016). The transcriptional auxin response is complex; numerous genes are controlled by auxin and are differentially expressed across tissues and organs depending on the developmental contexts (Kieffer et al., 2010; Salehin et al., 2015). Transcriptional auxin responses are mediated by the canonical TIR1/AFBs-Aux/IAA-ARF nuclear signaling pathway, which allows changes at the level of RNA in response to auxin in a timeframe of 3 to 5 minutes (McClure et al., 1989; Abel and Theologis, 1996) followed by changes in protein levels and activities at later time points. However, some auxin activities depending on the TIR1/AFB receptors, such as root growth inhibition (Fendrych et al., 2018) and root hair elongation (Dindas et al., 2018) are too fast to involve transcriptional regulation. This implies that TIR1/AFB auxin receptors, besides their well-characterized mechanism of transcriptional regulation, also trigger a so far unknown non-transcriptional signaling branch (Kubeš and Napier, 2019; Gallei et al., 2020).

In decades of auxin research, repeated observations of rapid cellular auxin effects have been made, including plasma membrane (PM) hyperpolarization, H^+^-pump activation, cytosolic Ca^2+^ or pH changes and protoplast swelling (Kubeš and Napier 2019). The underlying molecular mechanisms remain a mystery, but due to their rapidness, they were never linked to the canonical TIR1/AFB-Aux/IAA pathway and its downstream transcriptional responses. Prominent among those is the auxin regulation of clathrin-mediated endocytic trafficking of PIN auxin transporters (Adamowski and Friml, 2015; Narasimhan et al., 2020). This effect does not require the TIR1/AFB pathway and is presumably part of the feedback regulation of auxin on its intercellular auxin flow (Paciorek et al., 2005; Robert et al., 2010; Hajny et al., 2020). Moreover, a newly identified auxin analogue Pinstatic acid does not act via TIR1/AFB receptors, but still triggers distinct auxin responses, including the effect on PIN trafficking and auxin transport (Oochi et al., 2019). All these observations support existence of fast auxin responses mediated by so far unknown modes of auxin signaling and perception distinct from the well-studied nuclear TIR1/AFB mechanism (Gallei et al., 2020). Cell surface-localized TMK receptor-like kinases may contribute to this uncharacterized auxin signaling (Cao et al., 2019; Xu et al., 2014; Li, Verstraeten et al., 2021; Lin et al., 2021), but this remains unclear, mainly due to lack of an established auxin perception mechanism for this pathway.

Another key aspect of auxin action is directional auxin transport between cells (Vanneste and Friml, 2009). The establishment of auxin gradients and local maxima and minima in plant tissues can be attributed to local auxin biosynthesis (Morffy and Strader, 2020) and directional, intercellular auxin transport (Adamowski and Friml, 2015). Directionality of auxin flow is achieved by action and asymmetric (polarized) cellular localization of PIN auxin exporters at specific cell sides (Petrášek et al., 2006; Wiśniewska et al., 2006). PIN polarity and abundance at the PM depends on vesicle transport-related processes, such as constitutive recycling (Geldner et al., 2001) and clathrin-mediated endocytosis (Dhonukshe et al., 2007; Glanc et al., 2018), *de novo* secretion (Jásik et al., 2016; Salanenka et al., 2018) and degradation in lytic vacuoles (Abas et al., 2006; Kleine-Vehn et al., 2008a). Additionally, PIN phosphorylation at different residues is a key mechanism for PIN polarity or activity regulation (Michniewicz et al., 2007; Zhang et al., 2010; Barbosa et al., 2018; Tan et al., 2020, 2021; Xiao and Offringa, 2020). The cellular dynamics of the PIN-dependent auxin transport network have emerged as a prominent mechanism capable of translating endogenous (e.g. plant hormones) and exogenous (e.g. light and gravity) signals into developmental responses via relocating PIN transporters and thus redirecting routes of auxin transport (Vanneste and Friml, 2009; Adamowski and Friml, 2015).

Crucial regulation of PIN-mediated transport is by auxin itself. This positive feedback between auxin signaling and transport directionality is the key pre-requisite of the so-called auxin canalization (Sachs, 1981). The auxin canalization hypothesis proposes a mechanism for unique, self-organizing aspects of plant development such as the establishment of the apical/basal body axis during embryogenesis (Robert et al., 2013, 2018), flexible formation of vascular strands during organ formation (Benková et al., 2003; Heisler et al., 2005; Bhatia et al., 2016), shoot branching (Bennett et al., 2014; Zhang et al., 2020), leaf venation (Scarpella et al., 2006; Govindaraju et al., 2020), vasculature regeneration after wounding (Sauer et al., 2006; Mazur et al., 2016, 2020a; Hajný et al., 2020) and shoot gravitropic bending termination (Rakusová et al., 2016, 2019; Han et al., 2020). During canalization, auxin triggers coordinated PIN polarization to gradually establish directional auxin transport channels between auxin source and sink (Wabnik et al., 2010, 2011; Prát et al., 2018). Genetic and cell biological studies suggest that the auxin effect on PIN endocytic trafficking is central for PIN polarization and canalization (Sauer et al., 2006; Mazur et al., 2020a; Zhang et al., 2020). Nonetheless, auxin signaling mechanisms and downstream cellular processes, by which auxin regulates both PIN polarity and subcellular trafficking crucial for the self-organizing canalization processes, still remain elusive.

Here, we established a phospho-proteomics approach to identify components of the fast, non-transcriptional auxin responses. Among proteins that were rapidly phosphorylated in a TIR1/AFB-dependent and -independent manner, we further characterized the Myosin XI and MadB2 myosin binding proteins. We demonstrate that auxin-mediated phosphorylation of the myosin complex is essential for the feedback regulation of PIN auxin transporter trafficking and polarization, which is ultimately involved in canalization-mediated developmental processes such as vasculature formation in leaves or regeneration of wounded stems. Overall, our study uncovers novel downstream components of a so far elusive rapid auxin action and identifies Myosin XI proteins and their phospho-status as a crucial part of the mechanism, by which auxin regulates its own transport during plant development.

## Results

### Rapid auxin-mediated protein phosphorylation

Several responses to auxin occur within minutes or faster, and are too quick to be mediated by gene expression changes (Fendrych et al., 2018; Kubeš and Napier, 2019). Similar rapid responses such as for animal steroid hormones (Steinman and Trainor, 2010), or for osmotic stress in *Arabidopsis* (Stecker et al., 2014) have been attributed to changes in protein phosphorylation as prominent downstream signaling events.

Therefore, we tested whether auxin triggers rapid changes in protein phosphorylation and developed a procedure allowing to detect auxin-mediated phosphorylation events in times shorter than 2 minutes after auxin exposure. To maximize the detection depth for phospho-peptides in plant extracts, we compared a number of protein extraction and phospho-peptide enrichment strategies (**Figure S1A-J**). We found Filter-Aided Sample Preparation (FASP) to yield the best results, identifying most phospho-peptides with the least co-purification of nucleic acids (**Figure S1C**). We next combined FASP with various phospho-peptide enrichment strategies and obtained superior recovery with magnetic Ti^4+^ beads, identifying nearly 1500 phospho-peptides from Arabidopsis root extracts (**Figure S1A-B**). We found a significant overlap between the 3 phosphopeptide enrichment methods, but overall the Ti^4+^ beads resulted in the biggest portion of uniquely identified peptides (**Figure S1D**). When compared with the alternative enrichment methods, Ti^4+^ IMAC enrichment did not show a bias for charged peptides and mostly identified phospho-serines and -threonines (**Figure S1E**). TiO_2_(Mg) MOAC enrichment on the other hand resulted in more acidic phospho-peptides (**Figure S1E,I**). For the three evaluated methods, the overall nature of phosphorylated amino acids followed the previously published distribution: Serine(S) ~90%, Threonine(T) ~7 and Tyrosine(Y) ~1% (Wu et al., 2015) without discernable differences between them (**Figure S1F**). Moreover, also biochemical properties such as hydrophobicity or amino acid length did not differ significantly between the peptide enrichment methods (**Figure S1H-J**). In general, a protocol combining FASP and Ti^4+^ peptide enrichment was the best method for the identification of phospho-peptides in *Arabidospis thaliana* root samples, treated for 2 minutes with 100 nM indole-3-acetic acid (IAA) (**Figure 1A**).

**Figure 1.**
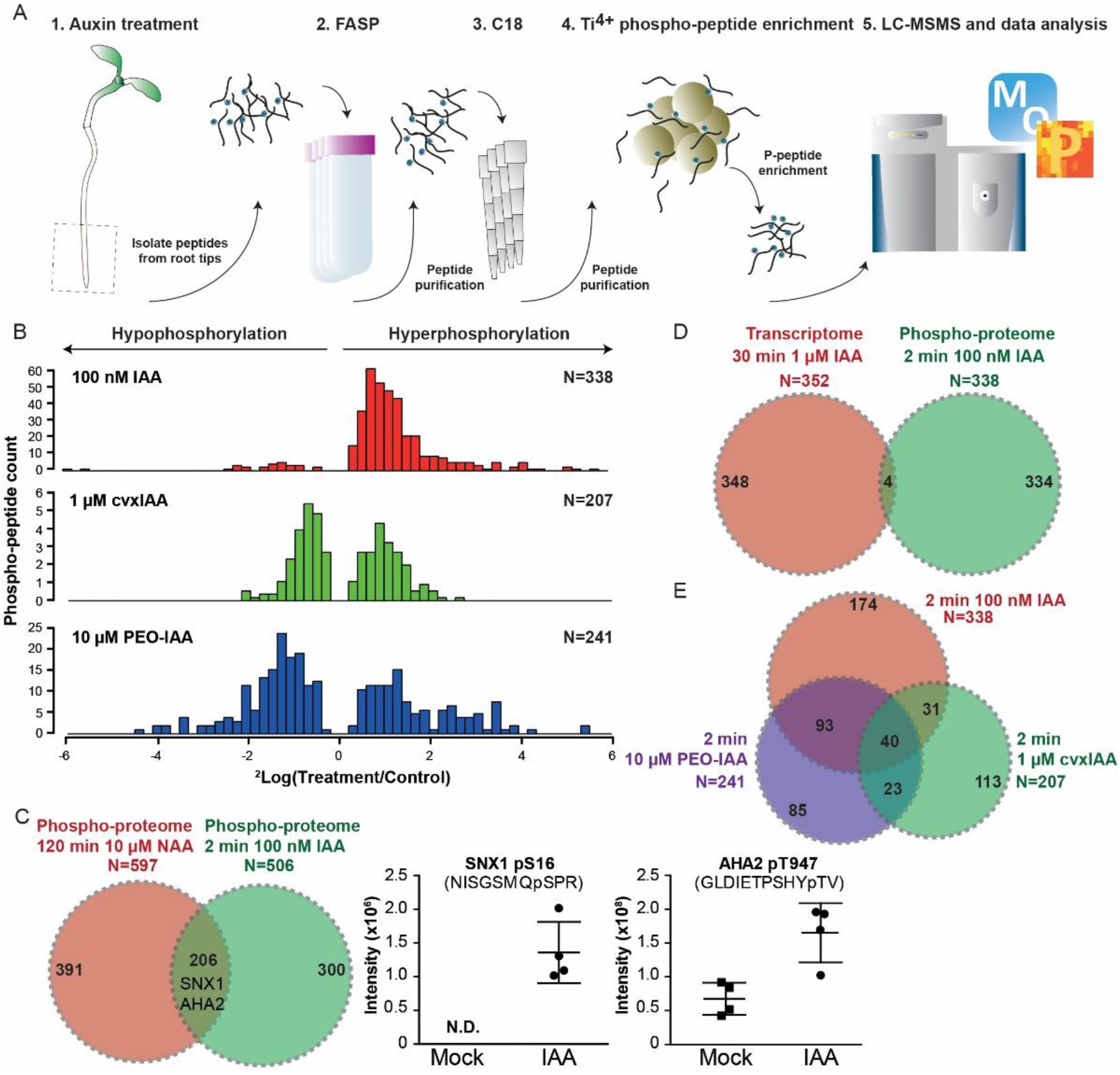
Phospho-proteomic analysis reveals rapid phosphorylation in response to auxin. (A) Schematic depiction of the phospho-peptide enrichment protocol. Root tips of 5 days old Arabidopsis seedlings were treated for 2 minutes and were harvested immediately. Proteins were extracted and digested to peptides that were submitted to the optimized FASP-C18-Ti^4+^ phospho-peptide enrichment protocol before analysis on LC-MSMS. (B) ^2^log fold changes of significantly hypo- and hyper-phosphorylated phospho-peptides (FDR ≤0.05) in WT treated for 2 min with either 100 nM IAA, 1 μM cvxIAA in cTIR/*ccvTIR1* or 10 μM PEO-IAA in WT compared to control treatment. (C) Comparison between the 2 min 100 nM IAA phospho-proteome dataset and a previously published phospho-proteome of roots stimulated with 10 μM NAA for 120 min (Zhang et al., 2013). The intersection contains specific auxin-upregulated phospho-peptides such as SNX1 and AHA2, for which MS intensity in the replicate samples of IAA and control treatments is depicted. (D) Comparison between differentially regulated phospho-peptides (FDR ≤0.05) in the short-term 100 nM IAA treatment and a published 30 min 1 μM IAA root transcriptome (Lewis et al. 2013). (E) Overlap of significantly regulated phospho-peptides by 100 nM IAA, 10 μM PEO-IAA and 1 μM cvxIAA treatment (FDR ≤0.05).

Our combined protein extraction and phospho-peptides enrichment method (**Figure 1A**) identified about 3100 phospho-peptides, which were subjected to a sensitive hybrid data analysis approach (Nikonorova et al., 2018) that allows detection of differential abundance across samples despite missing values. After filtering, the 2157 remaining phospho-peptides were subjected to FDR-controlled statistical comparison across treatments, resulting in about 10% differentially abundant phospho-peptides (FDR ≤ 0.05; **Figure 1B**). Global analysis showed that IAA mainly induced phosporylation, while limited auxin-dependent reduction in phosphorylation was detected (**Figure 1B**). The differential phospho-peptides map to 338 proteins (**Supplemental Table S1**) and among these are several proteins for which auxin-mediated phosphorylation has previously been shown (Zhang et al., 2013), such as SNX1 and AHA2 (**Figure 1C**). However, a large portion of the rapidly induced phospho-sites is unique (300/506; **Figure 1C**). Comparison of the proteins that are differentially phosphorylated with transcriptionally regulated genes (Lewis et al., 2013) confirmed that the early auxin-induced phospho-response targets a different and unique set of proteins (**Figure 1D**). Thus, auxin induces a rapid (within 2 minutes or shorter) protein phosphorylation response.

We next addressed whether the rapid phosphorylation response is mediated by the TIR1/AFB pathway using either PEO-IAA, an auxin antagonist for TIR1/AFB receptors (Hayashi et al., 2008) or the orthologous cvxIAA/ccvTIR1 system that specifically activates TIR1/AFB signaling (Uchida et al., 2018). Notably, cvxIAA in *ccvTIR1* roots induced some changes in phosphorylation, but the overlap with IAA-induced phosphorylation changes was limited (**Figure 1B, E**), suggesting that a substantial portion of the rapid auxin phospho-response was TIR1-independent (**Figure S2A**). PEO-IAA also rapidly induced changes in phosphorylation (**Figure 1B, E**), but again this phospho-response overlapped minimally with IAA-dependent or TIR1/AFB-dependent phospho-sites (**Figure 1E; Figure S2A-B**). The PEO-IAA treatment and the engineered cvxIAA/*ccvTIR* combination provide independent confirmation that the rapid phospho-response is, to a large extent, independent of the known TIR1/AFB auxin signaling.

Thus, we identified a novel, ultrafast auxin response that leads to differential phosphorylation of a range of proteins and is largely mediated through a yet unknown auxin perception mechanism.

### Auxin-mediated phosphorylation of Myosin XI and Myosin-binding proteins

The 338 proteins rapidly phosphorylated in response to auxin (**Supplemental Table S1**) reside in the nucleus, cytosol and at the PM (**Figure S2C**). The residues targeted by auxin-dependent phosphorylation were mostly serines, which follows the predicted phosphorylation pattern (**Figure S2D**; Wu et al., 2015). Among the rapidly phosphorylated proteins following auxin treatment were known regulators of auxin response such as 3-Phosphoinositide-Dependent protein Kinase 1 (PDK1), an AGC kinase that activates PIN auxin transporters (Tan et al., 2020; Xiao and Offringa, 2020), but the majority of the identified targets were novel. Some have reported roles in membrane trafficking (e.g. SYP132, BIG3, DRP2A/B, EPSIN2), microtubule regulation (e.g. MAP70-1, TOR1, NEK5) or chromatin biology (e.g. SUVR5, TPL, BRM, HDT1), but had not previously been associated with auxin-dependent processes. In short, the phospho-proteome dataset represents a rich starting point for exploring novel fast auxin responses.

Among the identified proteins (**Supplemental Table S1**), Myosin XIK (Uniprot code A0A1P8BCY2; encoded by AT5G20490), a motor protein with actin binding capability and presumably involved in endomembrane vesicular transport (Peremyslov et al., 2012), emerged as a prominent candidate phosphorylated in response to IAA treatment in several independent MS experiments (**Figure 2A; Supplemental Table S1**). In the globular tail domain (GTD) of Myosin XIK, the serine at position 1234 (S1234) was phosphorylated following IAA treatment, while phosphorylation of this site was not detected following PEO-IAA treatment and only minor differential phosphorylation was observed with cvxIAA treatment in *ccvTIR1* roots (**Figure 2A**), both suggesting largely TIR1/AFB-independent phosphorylation. We thus decided to independently confirm *in vivo*, auxin-mediated phosphorylation of the globular tail of Myosin XIK (S1234) by Western blot. To this end, we fused the GTD of myosin XIK with an RFP-tag and expressed the construct under the 35S promoter (*35S::MyosinXIK*^*WT*^ hereafter called *XIK*^*WT*^). This constructs were transformed into both, WT (Col-0) background and stable transformed seedlings were treated with 10 μM of synthetic auxin, 1-napththalene-acetic acid (NAA) for 30 minutes. Our Phos-tag Western blot analysis confirmed that Myosin XIK was more phosphorylated following auxin treatment (**Figure 2C**).

**Figure 2.**
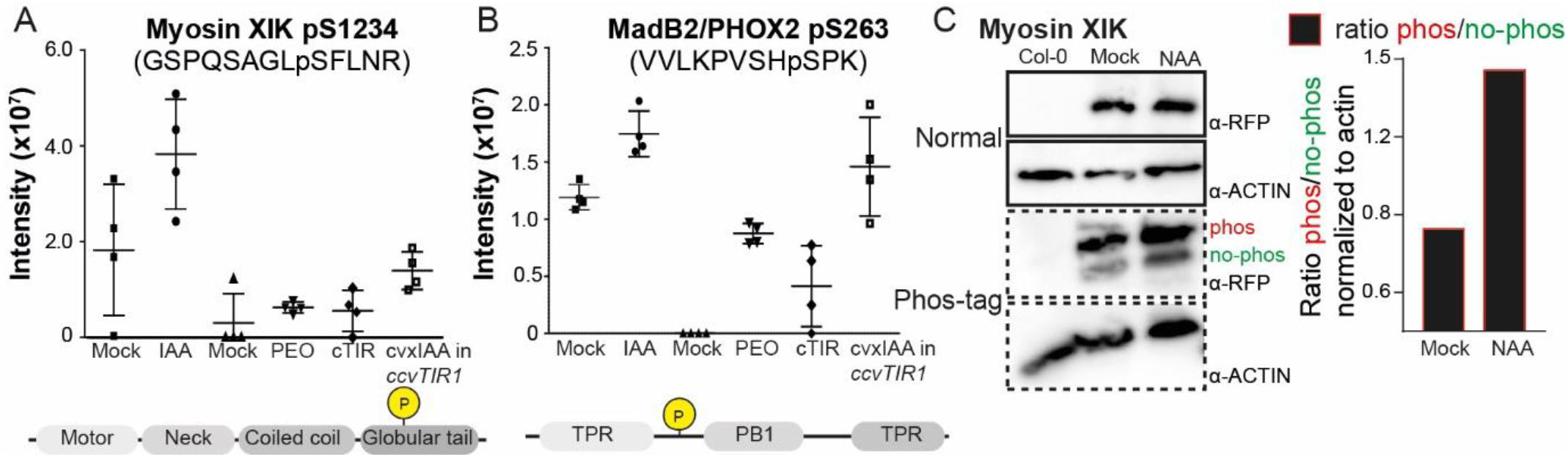
Rapid auxin-induced phosphorylation of Myosin XI and MadB2. (A) Relative MS intensity of the auxin-induced Myosin XIK phospho-peptide (containing S1234 as the significantly identified phospho-site) in 100 nM IAA, 10 μM PEO-IAA and 1 μM cvxIAA treated roots, always next to the respective mock (or control line such as cTIR1 for the cxvIAA treatment in the engineered *ccvTIR1* roots). In the schematic domain structure of Myosin XIK, the S1234 phospho-site located in the globular tail is indicated. Experiment performed in 4 independent biological replicates, represented as mean ± SD. (B) Relative MS intensity of auxin-induced phospho-peptide of MadB2/PHOX2 (containing S263 as the significantly identified phospho-site) in 100 nM IAA, 10 μM PEO-IAA and 1 μM cvxIAA treated roots, always next to the respective mock control (or control line such as cTIR1 for the cxvIAA treatment in the engineered *ccvTIR1* roots). In the schematic representation of MadB2 the S263 phospho-site is indicated. Experiment performed in 4 independent biological replicates, represented as mean ± SD. (C) Phosphorylation of Myosin XIK after auxin treatment visualized by Western and Phos-tag blot assays. 7 days old whole *35S::MyosinXIK^WT^* seedlings were mock or 10 μM NAA treated for 30 minutes after which proteins were extracted and submitted to Western and Phos-tag blot analysis. Quantification of the intensities of the target proteins in the Phos-tag blot compared to normal Western, normalized to the anti-actin loading control within the same sample lane, indicate a different phospho-state of the protein, which increased in the NAA-treated samples.

The *Arabidopsis* genome encodes 17 Myosin family members that are highly conserved (Reddy et al., 2001; Avisar et al., 2009; Ryan et al., 2017). Myosins have been shown to play roles in vesicle trafficking and endocytosis (Prokhnevsky et al., 2008; Peremyslov et al., 2012), gravitropism (Okamoto et al., 2015; Talts et al., 2016), auxin transport (Abu-Abied et al., 2018) and response (Ojangu et al., 2018). Protein alignment revealed that the S1234 phospho-site is conserved in Myosin XI proteins (**Figure S3A**). Despite not detected in the phospho-dataset, disruption of the homologous Myosin XIF, which is highly expressed in roots, results in similar trafficking defects (Avisar et al., 2008; Haraguchi et al., 2018). Therefore, we focused on both these close homologues, Myosin XIF and XIF.

Furthermore, we found Myosin binding protein MadB2/CLMP1/PHOX2 (AT1G62390) to be hyperphosphorylated following auxin treatment (**Figure 2B**). MadB2 has been shown to bind to Myosin XIK, but its function in plant development is less characterized (Kurth et al., 2017).

Thus, auxin induces rapid phosphorylation of Myosin XIK and the associated MadB2. This suggests that the identified ultrafast rapid auxin signaling targets myosin and associated proteins, thereby providing a potential mechanism to rapidly regulate cellular processes including auxin transport.

### Myosins XI in auxin-sensitive endomembrane PIN trafficking

Dynamic polar localization and thus action of PIN auxin transporters is tightly regulated by constitutive endocytic trafficking between PM and endosomes, which is actin-dependent (Geldner et al., 2001; Kleine-Vehn et al., 2008b) and thus potentially linked with myosin motor proteins function (Peremyslov et al., 2012). Furthermore, auxins, chiefly NAA, interfere with this PIN endocytic recycling providing a possible mechanism for feedback regulation of auxin transport (Narasimhan et al., 2020; Wabnik et al., 2011). We hence tested the involvement of Myosin XI proteins and their auxin-mediated phosphorylation in endocytic trafficking of PIN proteins, in particular.

Constitutive endocytic recycling of PIN proteins can be indirectly visualized by intracellular aggregation following treatment with the trafficking inhibitor Brefeldin A (BFA) (Geldner et al., 2001). After BFA treatment, the *myosin xik xif* double mutant displayed less intracellular PIN1 aggregation into so-called BFA bodies than the wild type (WT) (**Figure 3A-B**) implying defective PIN1 endocytic trafficking in the *myosin xik xif* mutant. Whereas in WT roots, NAA significantly reduced BFA body formation (Paciorek et al., 2005), we did not observe any such effects in *myosin xik xif* roots (**Figure 3A-B**).

**Figure 3.**
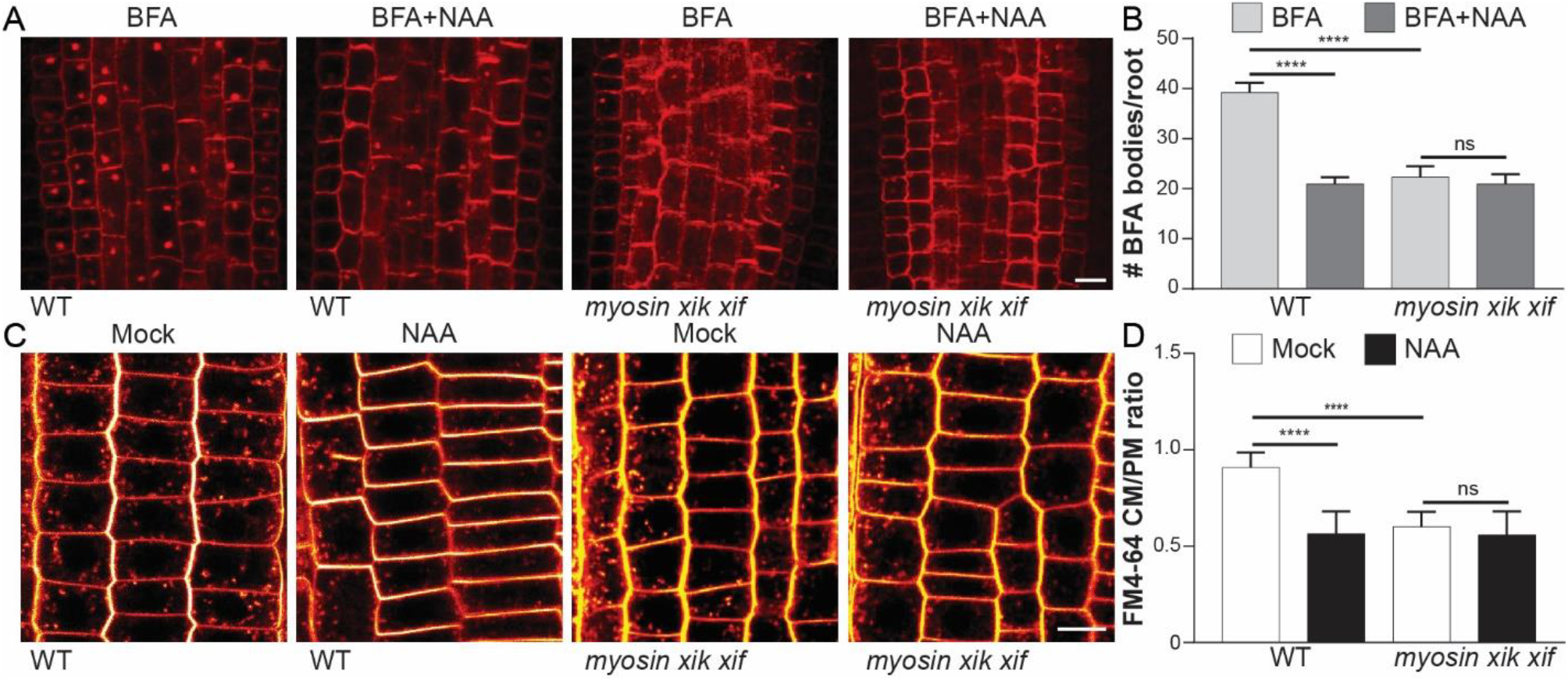
Myosin XI is involved in auxin-sensitive endomembrane PIN trafficking. (A) Representative images demonstrating the effect of BFA treatment or the co-treatment BFA + NAA in WT and *myosin xik xif* mutant roots. 3 days old seedlings were pre-treated with 10 μM NAA or DMSO mock for 30 minutes and then co-treated with 50 μM BFA and 10 μM NAA for another 60 minutes followed by anti-PIN1 antibody immunostaining according to Sauer et al. (2016). Scale bar = 10 μm. (B) Quantification of 50 μM BFA-induced bodies in WT and *myosin xik xif* roots and co-treatment of 50 μM BFA with 10 μM NAA. The average number of BFA bodies was determined per root, including a correction for the total number of cells evaluated per root. n > 20. Average # of BFA bodies per root ± SD is represented. **** for p ≤ 0.0001 and ns for p >0.05 determined by Tukey’s multiple comparison test. (C) Representative confocal images of FM4-64 staining in WT and *myosin xik xif* roots. PM staining and internalization of FM4-64 staining was quantified in 4 days old seedlings treated with 10 μM NAA or DMSO mock for 30 minutes. Scale bar = 10 μm. (D) Quantification of FM4-64 staining in WT and *myosin xik xif* roots. The CM/PM ratio was calculated by dividing the intercellular signal intensity by the PM-intensity. Bars represent the mean ± SD. Observations were made in more than 10 different roots and more than 20 cells per root were quantified. *** for p ≤ 0.001 and **** for p ≤0.0001 determined by Tukey’s multiple comparison test.

To asses a role of Myosin XI proteins in general trafficking, we followed the uptake of the lipophilic FM4-64 dye, which incorporates in the PM and gets into the cell by internalization to gradually mark endomembranes (Jelínková et al., 2010). Quantification of cytoplasmic (CM) FM4-64 staining and the PM-retained signal, revealed that FM4-64 uptake is significantly reduced in *myosin xik xif* mutants compared to WT (**Figure 3C-D**). Moreover, whereas NAA-treatment strongly reduced the internalization of FM4-64 in WT roots, we did not observe any additional effects of NAA in *myosin xik xif* roots (**Figure 3C-D**).

These observations demonstrate that Myosins XI are required for endomembrane trafficking and more specifically for the PIN1 endocytic recycling. Furthermore, the auxin-insensitivity of the *myosin* mutants in terms of PIN trafficking and global endocytosis indicates that the Myosin XI function may be required for the auxin-regulation of these trafficking processes.

### Auxin-mediated Myosin XI phosphorylation in endomembrane and PIN trafficking

Involvement of Myosin XI in auxin regulation of trafficking and its rapid auxin-triggered phosphorylation (**Figure 2**) prompted us to test the physiological relevance of the S1234 or S1256 phosphorylation in Myosin XIK and Myosin XIF,dv respectively. In the globular tail sequence, we mutated the respective Serine to Alanine to prevent phosphorylation (*MyosinXIK*^*S1234A*^ and *MyosinXIF*^*S1256A*^). Mutated sequences were C-terminally fused with an RFP-encoding tag, expressed under a *35S* promoter and introduced into WT (Col-0) and *myosin xik xif* mutant plants. Homozygous seedlings expressing the phospho-deficient Myosin XIK and XIF constructs (in Col-0 background) were treated with 10 μM NAA for 30 minutes to verify the phosphorylation status of Myosin XIK and XIF proteins in these mutants. Phos-tag and normal Western blot analysis showed that the phospho-deficient mutants were not phosphorylated following auxin treatment (**Figure S4A-C**).

We next examined whether the phosphorylation status of Myosin XIK and XIF contributes to regulation of PIN trafficking. When introduced into WT plants, the phospho-deficient mutants (*35S::MyosinXIK*^S1234A^ and *35S::MyosinXIF*^S1256A^, hereafter *XIK*^*S1234A*^ and *XIF*^*S1256A*^) showed less intracellular PIN1 aggregation in BFA bodies (**Figure 4A-B; Figure S4D**), less FM4-64 uptake (**Figure 4C-D; Figure S4E**) and were also less sensitive to auxin treatment (**Figure 4A-D; Figure S4D,E**). When introduced into the *myosin xik xif* mutant, the *XIK*^*WT*^ and *XIF*^*WT*^ constructs complemented the defective PIN1 accumulation in the BFA bodies (**Figure S5A-C**) and FM4-64 uptake (**Figure S5D-F**), whereas the phospho-deficient Myosin XI constructs were unable to rescue these defects (**Figure S5**).

**Figure 4.**
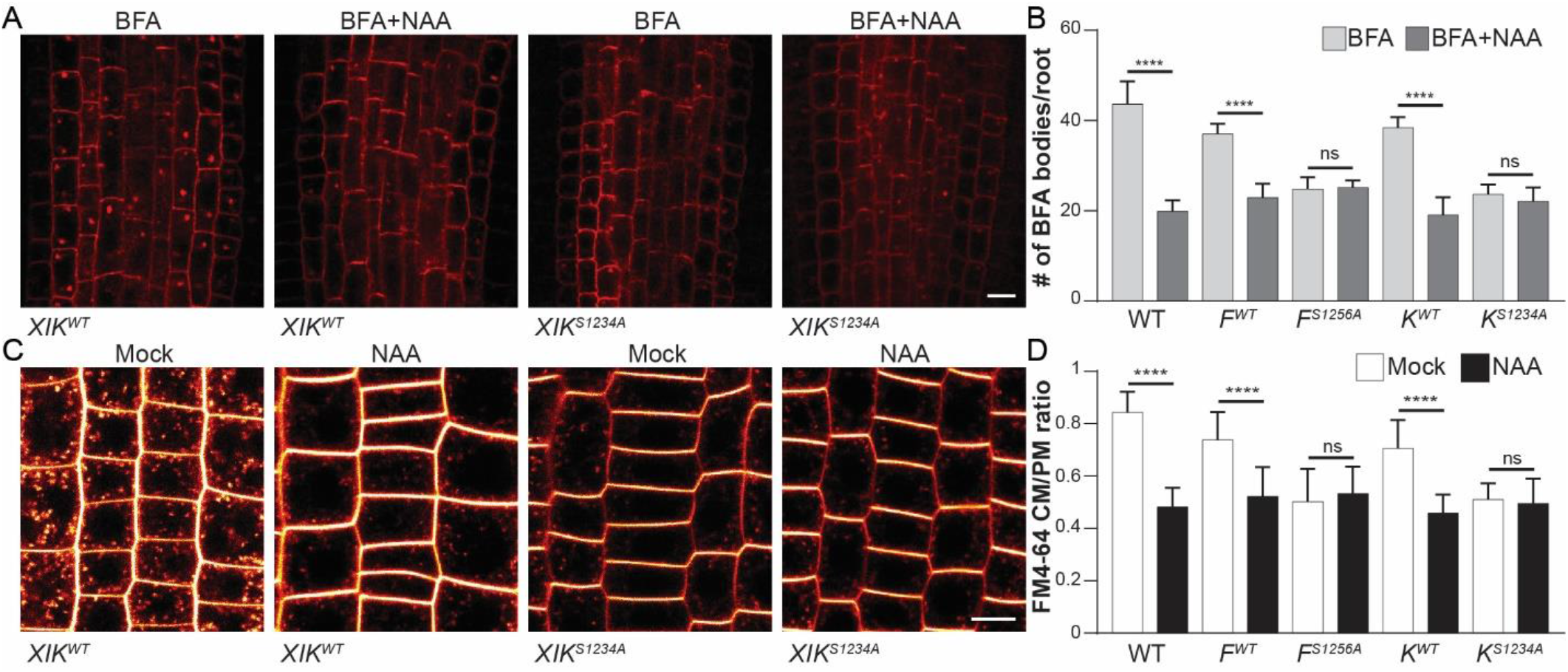
Auxin-mediated Myosin XI protein phosphorylation is required for PIN trafficking. (A) Representative set of images showing the effect of 50 μM BFA or 50 μM BFA + 10 μM NAA co-treatment in Myosin XIK^WT^ and phospho-deficient mutants (XIK^S1234A^) in WT background. Seedlings were immunostained using anti-PIN1 antibody according to Sauer et al. (2006). Scale bar = 10 μm. (B) Quantification of the average number of BFA bodies per root in the Myosin XI non-mutated and phospho-deficient lines in WT background. Data and error bars represent the mean ± SD. n > 15. **** p ≤ 0.0001 determined by Tukey’s multiple comparison test. (C) Representative confocal images of FM4-64 staining in *Myosin XIK*^*WT*^ and phospho-deficient mutants (*XIK*^*S1234A*^) in WT background. CM and PM staining were observed in 4 days old seedlings treated with 10 μM NAA or DMSO mock for 30 minutes followed by a staining using 2 μM of FM4-64. Scale bar = 10 μm. (D) Quantification of FM4-64 uptake in non-mutated and phospho-deficient Myosin XIF/XIK lines in WT background. The CM/PM ratio was calculated by dividing the internal staining intensity by the PM-intensity. Bars represent the mean ± SD of at least 40 cells in at least 10 independent roots. **** p ≤0.0001 determined by Tukey’s multiple comparison test.

Taken together, our results revealed that auxin-mediated phosphorylation of serine residues in the globular tail of Myosin XI proteins is critical for their function in auxin-sensitive endomembrane and PIN trafficking.

### Auxin-mediated Myosin XI phosphorylation in PIN1 repolarization in roots

The physiological relevance of auxin regulation of PIN endocytic trafficking is unclear. It has been proposed to be a key part of the cellular mechanism for auxin feedback on auxin transport and its directionality. This, in turn, is an essential pre-requisite of auxin canalization, a process allowing for the self-organized formation of polarized auxin channels during many developmental processes (Wabnik et al., 2010, 2011). Indeed, genetic and pharmacological manipulations revealed the crucial importance of auxin-regulated trafficking for auxin effect on PIN polarity and for formation of auxin transporting channels (Mazur et al., 2020a; Zhang et al., 2020; Hajný et al., 2020).

Therefore, we tested the involvement of Myosin XI in the auxin effect on PIN polarity, which can be visualized by PIN1 repolarization in auxin-treated roots (Sauer et al., 2006; Prát et al., 2018). As seen before, auxin induced a relocalization of PIN1 from the basal to the inner lateral side of endodermal cells in WT roots (**Figure 5A-C**) but, importantly, this auxin effect was not observed in the *myosin xik xif* mutant roots (**Figure 5A-C**). These observations demonstrate importance of Myosin XI for auxin-regulation of PIN polarity.

**Figure 5.**
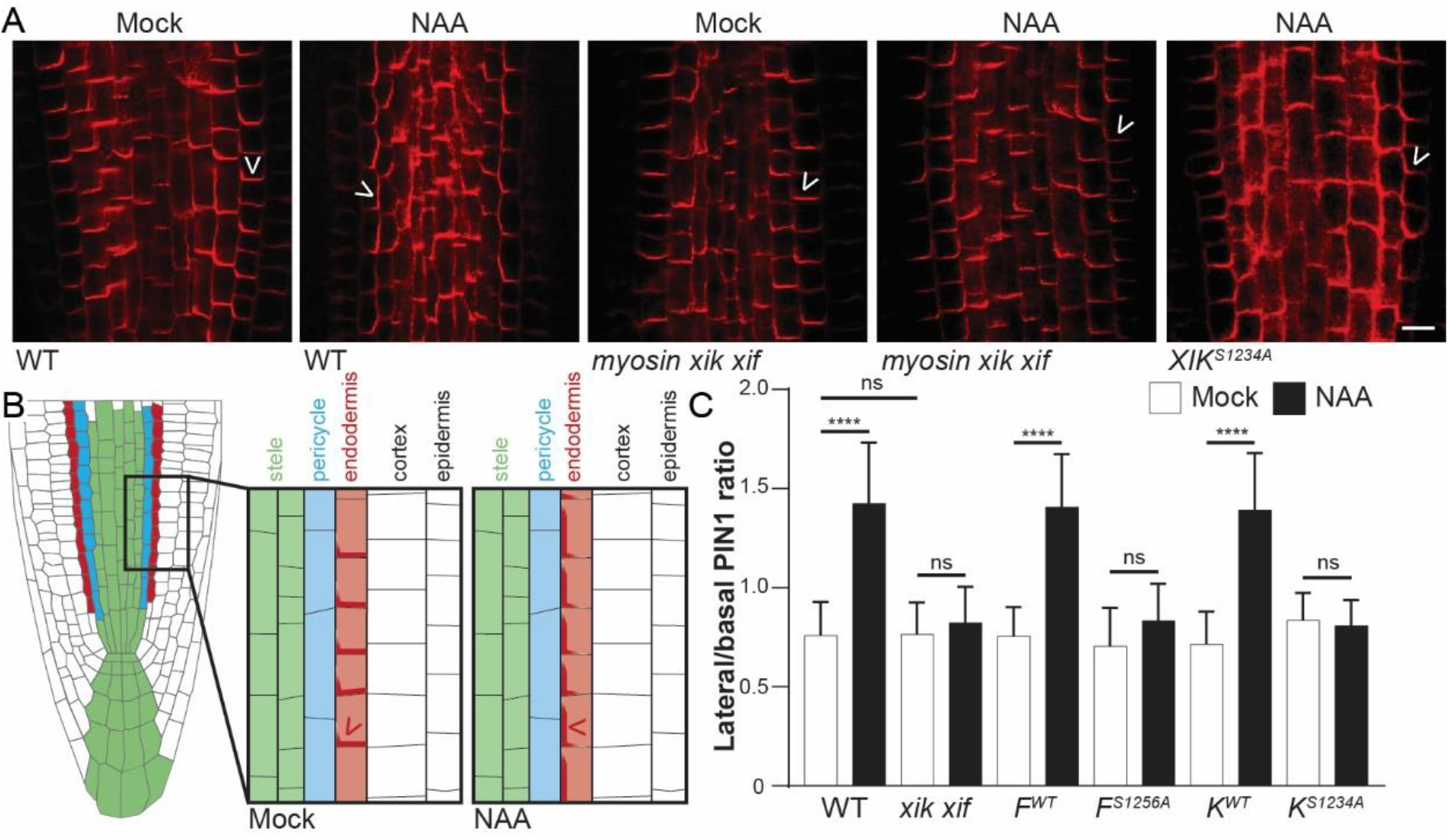
Auxin-mediated Myosin XI phosphorylation is required for PIN1 polarity regulation. (A) Representative images of DMSO mock and 10 μM NAA-induced PIN1 lateralization in WT, *myosin xik xif* mutants and roots expressing phospho-deficient *XIK*^*S1234A*^. The arrowheads indicate the predominant PIN1 localization in the endodermal cells. Scale bar = 10 μm. (B) Schematic representation of auxin-induced PIN1 lateralization in the endodermis of the root meristem. (C) Quantification of auxin-mediated PIN1 lateralization in endodermal root cells of WT, *myosin xik xif* and different lines expressing non-mutated or phospho-deficient *MyosinXIK/F* constructs (**Figure 5A** and **Figure S6A**). To visualize auxin-induced lateralization, the intensity ratio of lateral to basal PIN1 signal intensity was calculated per cell. Data represent mean ± SD. More than 50 cells were evaluated in at least 15 independent roots. **** p ≤0.0001 determined by Tukey’s multiple comparison test.

We then tested the ability of auxin to promote PIN1 repolarization in Myosin XI phospho-deficient mutants (in WT background). Our results showed that the phospho-deficient mutants were auxin-insensitive (**Figure 5A-C, Figure S6A**). Additionally, the defective auxin-mediated PIN1 repolarization in *myosin xik xif* mutants was rescued by introducing the non-mutated constructs (*XIF*^*WT*^ and *XIK*^*WT*^), but not by the phospho-deficient constructs (*XIF*^*S1256A*^ and *XIK*^*S1234A*^) (**Figure S6B-D**).

These results show that the Myosin XI function is required for auxin-mediated PIN repolarization in roots and that auxin-mediated phosphorylation of the serine residue in its globular tail is critical for the function in mediating auxin effect on PIN1 polarity.

### MadB2 myosin binding proteins in auxin-mediated regulation of trafficking and PIN polarity

Auxin induced phosphorylation not only of Myosin XI, but also of MadB2 Myosin binding proteins (**Figure 2B; Supplemental Table S1**). MadB2 has been shown to directly interact with Myosin XIK and it redundantly acts with its three paralogs in root hair growth (Kurth et al., 2017).

We examined whether MadB2 function is required for regulation of PIN polarity and trafficking. Following BFA treatment, *madb2* quadruple mutant (*madb 4KO*) accumulated less PIN1 in BFA bodies and also showed decreased sensitivity to auxin treatment (**Figures S7A**). Quantification of FM4-64 uptake demonstrated that endomembrane trafficking was also impaired in *madb 4KO* mutants and auxin treatment showed no additional inhibitory effect (**Figures S7B**). We also tested auxin-mediated PIN1 lateralization in *madb 4KO* mutant roots. Again, auxin failed to repolarize PIN1 to the lateral side of endodermal cells in *madb 4KO* roots (**Figure S7C**).

In conclusion, loss-of-function mutants in Myosin XI and in MadB2 Myosin binding proteins show essentially identical defects in auxin-regulated PIN trafficking and polarity. This supports that there is a common role for these proteins and their auxin-mediated phosphorylation in auxin feedback on PIN-dependent auxin transport.

### Auxin-mediated Myosin XI phosphorylation in PIN3 repolarization during shoot gravitropic bending termination

Next, we tested the relevance of Myosin XI and its auxin-mediated phosphorylation in different developmental contexts. During shoot gravitropic responses, gravity initially induces relocation of PIN3 to the bottom sides of endodermis cells to redirect auxin towards the lower side of the hypocotyl, which leads to growth promotion and upward hypocotyl bending. Subsequently, the auxin build-up at the lower side mediates a second relocation of PIN3 transporters to the inner lateral sides, thus re-establishing the symmetry of auxin fluxes and terminating asymmetric growth (Rakusová et al., 2016). Thus, the auxin effect on PIN3 polarity for the termination of shoot gravitropic bending represents an example of auxin feedback regulation of PIN-dependent auxin transport.

Indeed, the *myosin xik xif* mutant showed impaired polar auxin transport in hypocotyl (**Figure S8B**) and its hypocotyls were hyperbending (**Figure 6A-C**; Okamoto et al., 2015). Steady-state PIN3 polarity was not affected in *myosin xik xif* mutant hypocotyls (**Figure 6D**), neither was the gravity-induced PIN3 polarization in the endodermis cells after 6 hours gravistimulation (**Figure S9**). In contrast, the auxin-mediated re-establishment of PIN3 polarity at later stages (24 hours) was defective (**Figure S9**). To directly test the auxin feedback on PIN3 polarity in hypocotyls, we applied auxin exogenously, which induces a similar PIN3 repolarization to inner side of endodermal cells as observed at later stage of gravitropic response (Rakusová et al., 2016). Indeed, we observed normal auxin-induced PIN3 repolarization in WT hypocotyls, but not in *myosin xik xif* mutant hypocotyls (**Figure 6D; Figure S8A**). Thus, Myosin XI function is required, specifically, for the auxin-mediated PIN3 repolarization and for the gravitropic bending termination in hypocotyls.

**Figure 6.**
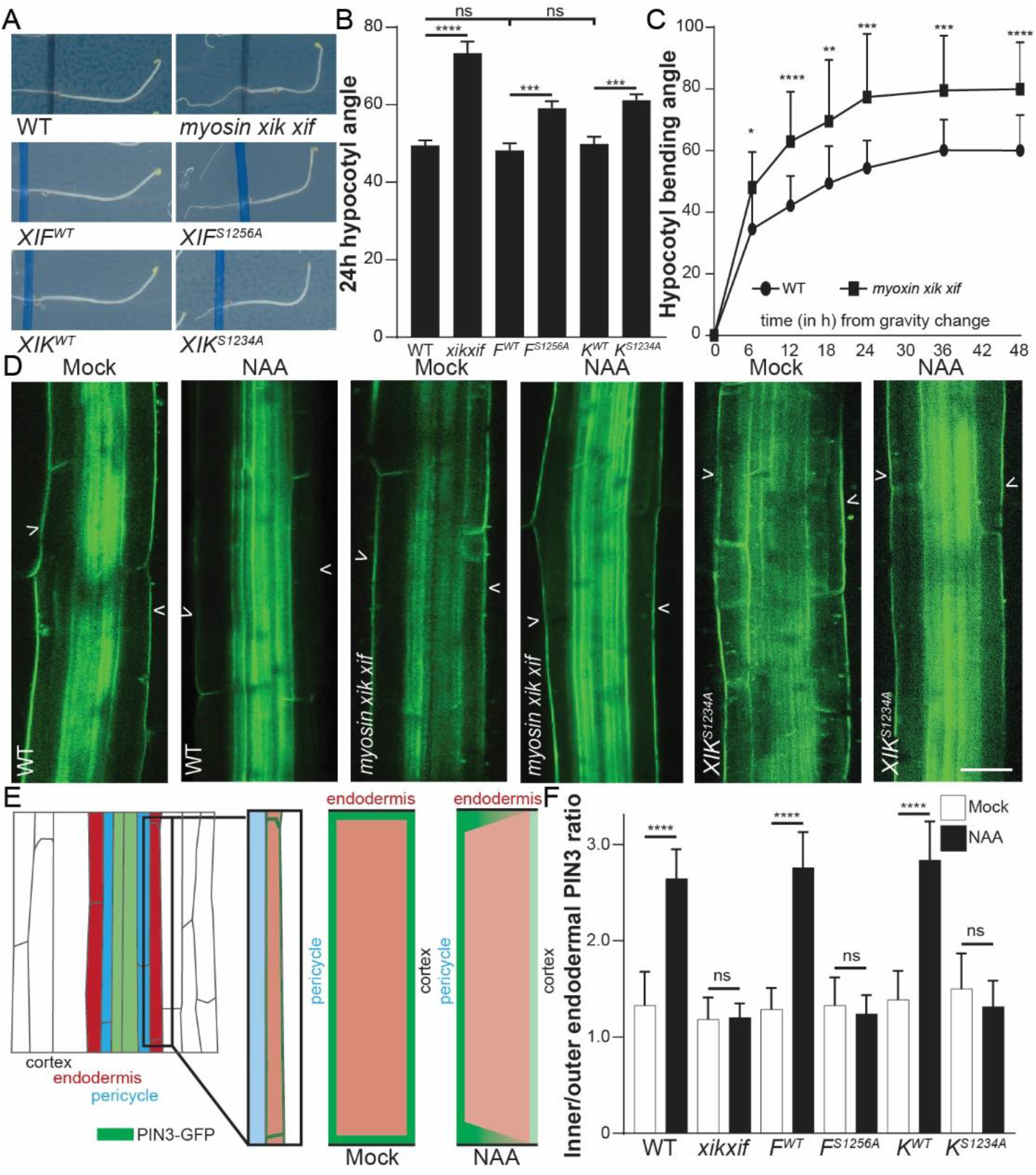
Phosphorylation of myosin XI proteins contributes to auxin-mediated PIN3 polarity regulation to terminate hypocotyl bending. (A) Representative seedlings showing gravity-induced hypocotyl bending. *myosin xik xif* bended stronger than Col-0 WT and similarly, the phospho-deficient Myosin XIF^S1256A^ and XIK^S1234A^ were bending more than their respective WT controls. (B) Bending angle of hypocotyls expressing Col-0 WT, *myosin xik xif,* non-mutated and phospho-deficient Myosin XIF and XIK constructs (in WT background). The bending angle was quantified in 3 days old etiolated seedlings gravistimulated for 24 hours. The myosin *xik xif* mutant and the phospho-deficient XIF^S1256A^ and XIK^S1234A^ showed hyperbending. (C) Kinetics of the gravitropic hypocotyl bending in WT and *myosin xik xif* seedlings. More than 30 hypocotyls were followed over time and the average bending angle ± SD is represented. ** p ≤ 0.01, *** p ≤0.001 and **** p ≤0.0001 determined by Student t-tests compared to the bending angle in WT at the same timepoint. (D) Representative images showing PIN3-GFP localization in WT, *myosin xik xif* mutant and hypocotyls of the phospho-deficient *XIK*^*S1234A*^ mutant (WT background) upon mock and 10 μM NAA treatment. Scale bar = 25 μm. (E) Schematic depiction of auxin-induced PIN3 repolarization in the endodermis of WT etiolated hypocotyls. (F) Quantification of auxin-mediated PIN3 repolarization in the hypocotyl of WT, *myosin xik xif* and the different *XIF*^*WT*^/*XIK*^*WT*^, and phospho-deficient mutants (in WT background) (**Figure 6A** and **Figure S8A**). The inner/outer endodermal ratio is determined by dividing the PIN3-GFP fluorescence signal intensity between inner side and outer side of hypocotyl endodermal cells. Data and error bars represent the mean ± SD.

We also tested whether the auxin-regulated Myosin XI phospho-sites is essential for PIN3 polarization. Gravity-induced PIN3 polarization was normal in both phospho-deficient (*XIF*^*S1256A*^; *XIK*^*S1234A*^) and non-mutated control myosin mutants (**Figure S9**). On the other hand, auxin-mediated PIN3 repolarization at later stages of gravity response (**Figure S9B-C**), or after exogenous auxin application (**Figure 6D; Figure S8A**), was defective in the phospho-deficient mutants, whereas the non-mutated controls showed normal PIN3 repolarization. Accordingly, the hypocotyls of Myosin XI phospho-deficient mutants were hyperbending (**Figure 6A-B**).

These observations demonstrate that Myosin XI function and its auxin-mediated phosphorylation are required for auxin-mediated PIN3 repolarization in shoot and for the resulting termination of the gravitropic bending.

### Auxin-mediated Myosin XI phosphorylation in auxin canalization-mediated vasculature formation and regeneration

The classical auxin canalization hypothesis proposes auxin feedback regulation of directional auxin transport as the key part underlying self-organizing aspects of plant development, such as vasculature formation during spontaneously arising leaf venation patterns (Scarpella et al., 2006) and flexible regeneration of vasculature around wounding sites (Sauer et al., 2006; Mazur et al., 2016, 2020a, b).

We first analyzed cotyledon venation patterns in *myosin xik xif* mutant seedlings. As compared to WT leaves, *myosin xik xif* mutants showed typically fewer loops (Figure S8E). We also analyzed seedlings overexpressing non-mutated and phospho-deficient versions of MyosinXI proteins. Cotyledons of lines expressing the *XIK*^*WT*^ and *XIF*^*WT*^ constructs did not show any pronounced defects in the number of loops (**Figure S8E**). On the other hand, seedlings expressing phospho-deficient constructs (*XIK*^*S1234A*^ and *XIF*^*S1256A*^) had a higher occurrence of fewer loops (**Figure S8E**). Thus, mutants defective either in Myosin XI function or in their auxin-mediated phosphorylation show mild defects in cotyledon venation.

Next, we analyzed vasculature regeneration in stems after wounding. As visualized by toluidine blue staining, within 7 days after wounding the vasculature fully developed in WT stems and reconnected around the interruption while, in *myosin xik xif* mutant stems, the newly formed vasculature was partially connected (**Figure 7A**). We also analyzed the regeneration in phospho-deficient Myosin XI transgenic plants (WT background)(**Figure 7B**). The Myosing XI (*XIK*^*W*T^ and *XIF*^*WT*^) stems showed partial regeneration around the wound, but the phospho-deficient Myosin XI stems (*XIK*^*S1234A*^ and *XIF*^*S1256A*^) did not regenerate at all (**Figure 7B**). Thus, mutants defective either in Myosin XI function or in their auxin-mediated phosphorylation show defects in vasculature regeneration around the wound.

**Figure 7.**
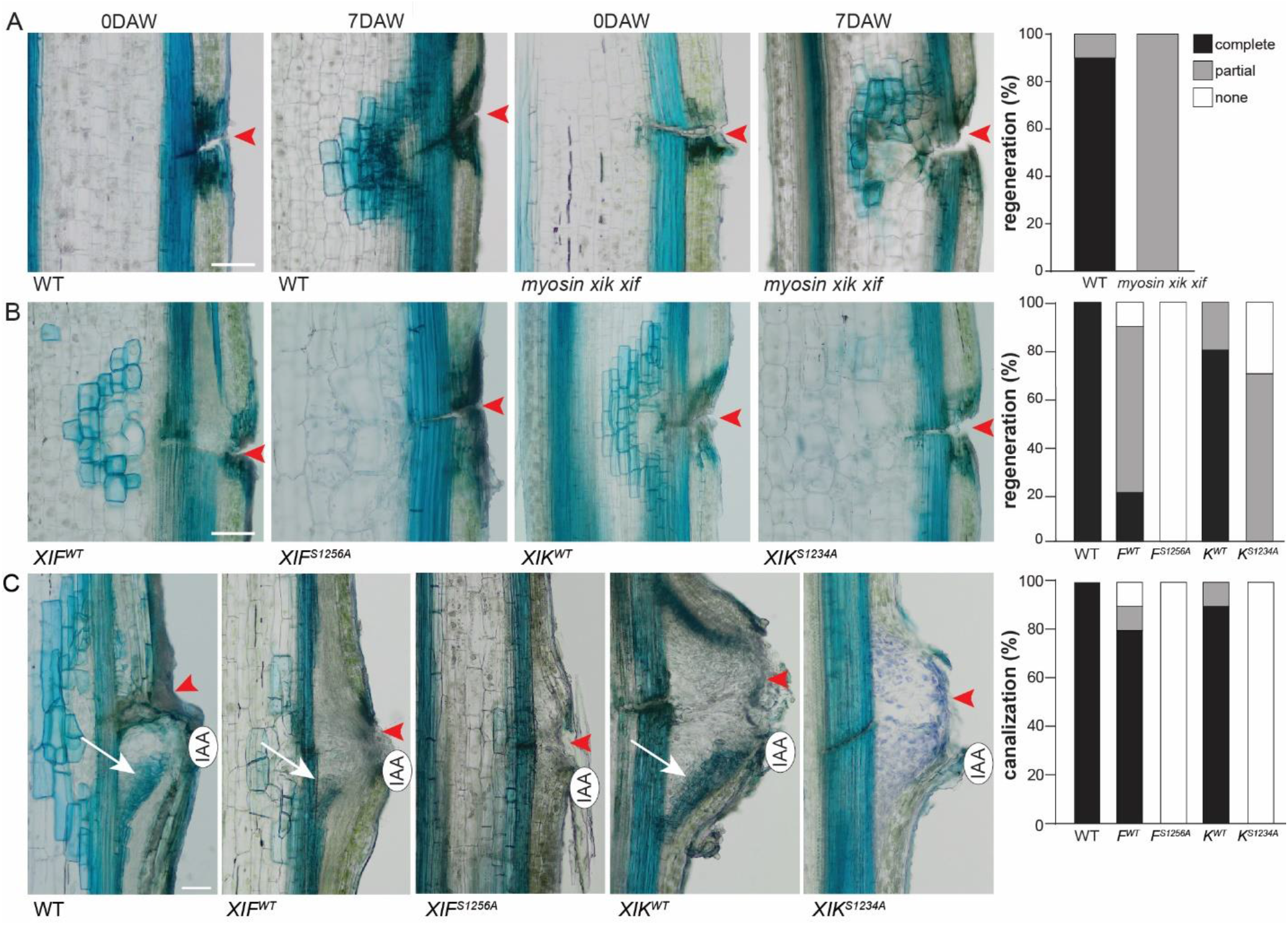
Auxin canalization-mediated vasculature formation and regeneration depends on auxin-mediated Myosin XI phosphorylation. (A) Vasculature regeneration in wounded *Arabidopsis thaliana* stems. The incision site is marked by a red arrow. 7 days after wounding (DAW) all WT stems regenerated vasculature tissue around the wound, visualized by toluidine blue staining while in the *myosin xik xif* mutants, this regeneration was defective and the newly formed vasculature was incomplete. Scale bar = 100 μm. (B) Representative sections and quantification of vasculature regeneration in wounded *Arabidopsis* stems of *XIF*^*WT*^/*XIK*^*WT*^ and the phospho-deficient mutants (WT background). Whereas the vasculature in stems expressing the WT constructs (XIF^WT^ and XIK^WT^) was regenerating, we did not observe vasculature regeneration in stems expressing the phospho-deficient constructs (XIF^S1256A^ and XIK^S1234A^). Red arrowhead shows the incision site. n =20. Scale bar = 200 μm. (C) Auxin-induced canalization in wounded *Arabidopsis* stems of XIF^WT^/XIK^WT^ and the phospho-deficient mutants (WT background). Exogenous IAA application (indicated by the oval shape) triggered the formation of a channel (indicated by the white arrow) from this source to the existing vascular tissue (visualized by toluidine blue). Channel formation was observed in stems expressing *XIF*^^WT^^ and *XIK*^*WT*^. However, regeneration failed in stems expressing the phospho-deficient constructs (*XIF*^*S1256A*^ and *XIK*^^S1234A^^). Red arrowhead shows the incision site and the white arrow indicates the newly formed channel. Scale bar = 200 μm.

The most direct manifestation of auxin-induced canalization processes, is the formation of a auxin-transporting channels followed by vasculature differentiation starting at local, exogenous auxin source (Mazur et al., 2016, 2020a, b). In WT stems a distinct, new vasculature strand connecting the external IAA drop to the existing vasculature was formed; however, in *myosin xik xif* mutant stems, no such formation of well defined vasculature was observed (**Figure 7C**). Similar, auxin application induced new vasculature formation in *XIF*^*WT*^ and *XIK*^*WT*^ stems, but not in phospho-deficient Myosin XI mutants stems (**Figure 7C**).

These results show that Myosin XI proteins and their auxin-mediated phosphorylation are important for the auxin canalization-mediated vasculature formation; to a limited extent during leaf venation but indispensable for the vasculature regeneration around a wound as well as for the *de novo* vascular strands formation from the external auxin source. This collectively shows that auxin-mediated myosin XI phosphorylation, which at the cellular level mediates auxin regulation of PIN trafficking and polarity, is required for developmental processes that depend on coordinated, auxin-regulated polarization of PIN-dependent auxin transport.

## Discussion

Auxin, a crucial regulator of plant development, is known to trigger well-characterized nuclear signaling to mediate transcriptional reprogramming, nonetheless the existence of rapid, non-transcriptional cellular auxin effects has been discussed for decades (Kubeš and Napier, 2019). Here, we established a fast phospho-proteomics approach, identify a thus far unknown rapid phosphorylation response, and characterize novel targets of this rapid auxin signaling pathway. Among the identified candidates that were phosphorylated within few minutes in response to auxin, we characterized in depth Myosin XI proteins and the MadB2 Myosin binding protein. This analysis identified their auxin-mediated phosphorylation as an essential part of the feedback mechanism, by which auxin regulates its directional transport for the self-organizing aspects of plant development.

### Auxin mediates rapid protein phosphorylation by distinct signaling pathways

Protein phosphorylation and de-phosphorylation are prominent post-translational modification to control protein functions and it is among the most common downstream events in many signaling pathways. Therefore, in order to identify new players of the rapid auxin responses, we optimized a phospho-proteomics protocol and designed treatment conditions in Arabidopsis seedling root tips, allowing identification of proteins phosphorylated within 2 minutes of auxin treatment with a high coverage. By using the available tools, such as the auxin antagonist specific for the TIR1/AFB receptors, PEO-IAA and the engineered receptor-ligand pair cvxIAA/*ccvTIR1*, we were able to distinguish auxin-induced modifications, which are dependent and independent of the canonical TIR1/AFB1 signaling.

To our knowledge, this rapid phosphorylation response is among the fastest responses to auxin reported to date. The set of proteins that is targeted by rapid auxin-dependent phosphorylation is distinct from those targeted by transcriptional control (**Figure 1D**), and thus represents a functionally unique aspect of auxin activity. The identified proteins targeted by this rapid response are diverse in nature and function, and provide a rich source of hypotheses on mechanisms mediating ultrafast, non-transcriptional auxin responses. They for example include chromatin and transcriptional regulators, mediators of membrane trafficking, regulators of RNA splicing and -suppression and various groups of enzymes. It is likely that these phosphorylation events have a functional impact on many of these pathways. Given that the response is, at least in part, independent of the TIR1 receptor, there is a potential for this response to occur in species that do not have a TIR1 orthologue, such as the *Charophyceae* algae (Mutte et al., 2018). Key future questions will therefore be, what receptor and downstream components mediate this ultrafast, non-transcriptional auxin response, and how universal it is in the plant kingdom.

### Auxin regulates PIN trafficking and polarity via phosphorylation of myosin complex

Myosin XIK and its interactor myosin-binding protein MadB2 were identified among the confirmed candidates identified as being rapidly phosphorylated by TIR1-independent auxin signaling. Given the concomitant auxin regulation of these group of functionally related proteins and the potential role of myosin in cellular processes related to trafficking and polarity, we focused on these candidates as potential new components of auxin feedback on its own polar, intercellular transport, which is the key prerequisite of so-called auxin canalization (Sachs, 1981).

The auxin canalization hypothesis provides a mechanistic framework for flexible, self-organizing plant developmental processes such as post-embryonic organogenesis, shoot branching, leaf venation and vascular tissue regeneration. A key condition for canalization is the feedback regulation of auxin transport, as manifested at a cellular level by the auxin effect on the polar, subcellular localization of PIN auxin transporters. Constitutive PIN trafficking and clathrin-mediated endocytosis represent key cellular components of this feedback regulation of PIN polarity (Paciorek et al., 2005; Robert et al., 2010; Mazur 2020a; Zhang et al., 2020).

Our analysis revealed that the Myosin XI proteins are required for auxin-regulated PIN trafficking and PIN polarity rearrangement in different developmental contexts. This is demonstrated by strong defects in the corresponding loss-of-function mutants, where auxin is largely unable to affect PIN subcellular trafficking or mediate PIN repolarization. Notably, similar defects in auxin-mediated PIN trafficking and polarization were also observed in the phospho-deficient Myosin XI transgenic lines, in which we mutated the identified auxin-dependent phospho-site. This shows that auxin-mediated phosphorylation of Myosin XI proteins is crucial for auxin-mediated PIN trafficking and polarity regulation.

The identification of a large number of myosin interacting proteins indicates there is a series of specific interactions that allow myosin motors to operate on various cargoes (Peremyslov et al., 2013; Kurth et al., 2017) that likely also include PIN proteins, which traffic along the actin cytoskeleton (Geldner et al., 2001). Our genetic analysis demonstrates that the MadB2 myosin binding protein family contributes to PIN trafficking and polarity regulation in roots. The most likely scenario is that PIN proteins are cargoes of vesicles that are decorated by the MadB2 family proteins, which bind to the Myosin XI proteins forming a transport network (Peremyslov et al., 2013; Kurth et al., 2017). By this mechanism, PIN proteins can be transported though the trafficking pathway to specific, polar destinations. Notably, auxin-mediated phosphorylation of these components provides a means how auxin itself can influence this process in different developmental contexts. The topics of future investigations include how the myosin binding proteins convey specificity for different cargoes, how myosin tail-binding proteins are employed by myosins to carry out their trafficking function and not least, how the auxin-mediated phosphorylation influences the whole system to aid the required PIN polarity re-arrangement to mediate redirection of auxin fluxes in different developmental processes.

### Auxin feedback on directional auxin transport in plant development

With identification of Myosin XI and corresponding myosin-binding proteins, we obtained molecular components of the feedback between auxin signaling and polarity re-arrangements of PIN auxin transport components. This provides a genetic means to test the auxin canalization hypothesis, its cellular requirements and developmental roles.

Firstly, the analysis of *myosin xik xif* and *madb 4KO* mutants confirmed the previously proposed link between auxin effect on PIN trafficking and auxin effect on PIN polarity (Sauer et al., 2006; Wabnik et al., 2010, 2011; Mazur 2020a; Zhang et al., 2020) because these mutants are defective in both processes and their auxin regulation. Moreover, they also allowed to link auxin effect on PIN polarity, typically visualized in roots, to similar auxin-mediated PIN rearrangements in other developmental contexts, mainly those involving flexible formation of auxin channels determining position of new vasculature, such as vasculature regeneration after wounding in stems. Additionally, our observations confirmed that the Myosin XI-based mechanism also operates in other developmental processes, which do not involve auxin channel formation, but where auxin effect on PIN polarity has been observed, such as PIN3 polarization for gravitropic bending termination in shoots. Hence, our findings provide mechanistic and molecular support for the auxin canalization hypothesis and show that the myosin transport network and its auxin regulation play an essential role in many developmental processes involving auxin canalization or auxin effect on its transport directionality.

In conclusion, this study demonstrates that Myosin XI and MadB2 myosin binding proteins, as part of the myosin transport network, are required for auxin feedback on PIN polarity, more specifically for auxin effect on PIN trafficking and polarity re-arrangements. Furthermore, we also uncover that auxin-mediated phosphorylation of these myosins is required and sufficient for their function in PIN trafficking and polarity regulation. Our work provides first mechanistic insights into feedback regulation of auxin transport and reveals its crucial role in many self-organizing developmental processes involving auxin canalization.

## Acknowledgments

We thank Prof. Ikuko Hara-Nishimura (Kyoto University, Japan) and Prof. Valerian V. Dolja (Oregon State University) for sharing published lines. We acknowledge Matyáš Fendrych for initiating the phospho-proteomics approach. This work was supported by the European Research Council (ERC ETAP: 742985) and the Austrian Science Fund (FWF, grant number I 3630-B25) to J.F., and by the Netherlands Organization for Scientific Research (NWO; VICI grant 865.14.001) to D.W. H.H. was supported by the China Scholarship Council (CSC scholarship).

## Author Contributions

H. H. and J. F. designed research. H. H. performed most of the experiments. I. V. synthesized the Myosin XIK and XIF GTD sequences and performed the Western blot assays. M. R. and D. W. optimized, performed and analyzed the phosphoproteomics experiments. J. H. carried out the auxin transport assay. K. Ö. set up the PhosTag method. E. M. and N. R. performed the regeneration assays. H. H., I.V. and J. F. wrote the manuscript with input from all other authors.

## Methods

### Plant material and growth

The following transgenic and mutant lines were used: WT (Col-0), *pPIN3::PIN3-GFP* (Žádníková et al., 2010), *myosin xik xif* (Okamoto et al., 2015), *madb 4KO* (Kurth et al., 2017). Mutant combinations in *PIN3::PIN3-GFP* were generated through genetic crosses. Seeds were surface-sterilized by chlorine gas, sown on half-strength Murashige and Skoog (½ MS) medium supplemented with 1% (w/v) sucrose and 0.8% (w/v) phyto agar (pH 5.9), stratified in the dark at 4°C for 2 days and then grown vertically at 21°C with a long-day photoperiod (16h light/8h dark). Light sources used were Philips GreenPower LED production modules [in deep red (660 nm)/far red (720 nm)/blue (455 nm) combination, Philips], with a photon density of 140.4 μmol/m2/s ± 3%.

### Protein extraction

Five days after germination, roots were treated with either 100 nM IAA, 10 μM PEO-IAA or 1 μM cvxIAA or their respective solvents as mock. Root tips were directly harvested on liquid nitrogen. For IAA and PEO-IAA treatments, Col-0 WT was used, while for cvxIAA treatment, *ccvTIR1* seeds were used (Uchida et al., 2018). The harvested root tips were ground to fine powder in liquid nitrogen and extracted in SDS lysis buffer (100mM Tris pH8.0, 4%SDS and 10mM DTT). Protein extract was sonicated using a cooled Biorupter (Diagenode) for 10 minutes using high power with 30 seconds on 30 seconds off cycle. The lysate was cleared by centrifugation at maximum speed (13,000xg) for 30 minutes. Protein concentrations were determined using Bradford reagent (Bio-Rad).

### Protein precipitation

Acetone precipitation was done according to Humphrey et.al. (2015). Methanol chloroform precipitation was done according to Vu et.al. (2016). For trichloroacetic acid (TCA) precipitation 1 volume of ≥99% TCA was added to 4 volumes of protein lysate. Mixtures were incubated on ice for 10 minutes and spun at maximum speed (13,000xg) for 5 minutes at 4◦C. Pellet was washed twice with acetone at maximum speed (13,000xg) for 5 minutes at 4◦C and then air dried and resuspended in 50 mM ammoniumbicarbonate (ABC) (Sigma).

### Filter aided sample preparation (FASP)

For FASP, 30 kDa cut-off amicon filter units (Merck Millipore) were used. Filters were first tested by applying 1000 μl urea buffer UT buffer (8 M Urea and 100mM Tris, pH 8.5) and centrifuging for 20 minutes at 6,000 RPM. All further centrifuging steps were at this speed and temperature. The desired amount of protein sample was next mixed with UT buffer to a volume of 5000 μl, applied to filter and centrifuged for 20 minutes. Filter were washed with UT buffer and centrifuged for 20 minutes. Retained proteins were alkylated with 50 mM acrylamide (Sigma) in UT buffer for 30 minutes at 20◦C while gently shaking. Filter was centrifuged and afterwards washed three times with UT buffer for 20 minutes. Next, filters were washed three times with 50 mM ABC buffer. After the last wash, proteins were cleaved by adding Trypsin (Roche) in a 1:100 trypsin:protein ratio. Digestion was completed overnight. Filter was transferred to a new tube and peptides were eluted by 20 minutes centrifugation. Further elution was completed by two times adding 50 mM ABC buffer and centrifuging.

### C18 Stage tip clean up

For peptide desalting and concentrating, 1000 μl tips were fitted with 2 plugs of C18 octadecyl 47mm Disks 2215 (Empore™) material and 10 *μg* of LiChroprep® RP-18 peptides (Merck). Tips were sequentially equilibrated with 100 % methanol, 80 % ACN in 0.1 % formic acid and twice with 0.1 % formic acid with intermitting centrifuging for 4 minutes at 1,500xg. After equilibration, peptides were loaded and centrifuged for 20 minutes at 400xg. Bound peptides were washed with 0.1 % formic acid and eluted with 80 % ACN in 0.1 % formic acid by spinning for 4 minutes at 1,500xg. Eluted peptides were subsequently concentrated using a vacuum concentrator for 30 minutes at 45◦C and resuspended in 50 μl 0.1 % formic acid.

### Phosphopeptide enrichment

For the magnetic Ti^4+-^IMAC (MagResyn) and TiO_2_-MOAC (MagResyn) approaches, the manufacture’s protocols were followed without modifications (Resyn Biosciences). For stage tip based TiO_2_ Titansphere™ (GL Sciences) a 1:2 peptide to TiO_2_ (μg/μg) was used. FASP eluted peptides were mixed with ACN and TFA to a concentration of 50 % ACN and 6 % TFA. TiO_2_ columns were made with double C8 membrane and desired amount of beads in 100 % methanol. The columns were washed and equilibrated with 100 % ACN and 80 % ACN in 6 % TFA using centrifugation for 4 minutes at 1,500xg. Sample was loaded at 400xg for 30 minutes. Non-specifically bound peptides were washed with 80 % ACN in 6 % TFA by centrifugation for 4 minutes at 1,500xg and 2 times with 60 % ACN in 1 % TFA for 4 minutes at 1,500xg. Next, bound phosphopeptides were eluted three times in 40 % ACN and 15 % NH4OH. After the last elution samples were concentrated using a vacuum concentrator for 30 minutes at 45◦C. Samples were subsequently acidified using 10% TFA and processed with C18 Stagetip clean up.

### Mass spectrometry and data analysis

Phosphopeptides were applied to online nanoLC-MS/MS using a 120 min acetonitrile gradient from 8-50 %. Spectra were recorded on a LTQ-XL mass spectrometer (Thermo Scientific) according to Wendrich et.al. (2017). Data analysis of obtained spectra was done in MaxQuant software package (Wendrich et al., 2017), with the addition of phosphorylation as a variable modification. Data analysis and visualization was performed in Perseus, Adobe Illustrator and R. Filtering of phosphopeptides was conducted using a hybrid data filtering approach as described previously (Nikonorova et al. 2018). Protein subcellular localisation analysis was preformed using the Multiple Marker Abundance Profiling (MMAP) tool on the subcellular localisation database for Arabidopsis proteins (SUBA4) (Hooper et.al. 2017).

### Myosin XIK and XIF phosphorylation mutagenesis

The Myosin XIK and XIF GTD sequences (from amino acid 1100 to the C terminus) (Peremyslov et al., 2008) with mutations S1234A (*XIK*^*S1234A*^), S1256A (*XIF*^*S1256A*^) or in non-mutated version (*XIK*^*WT*^ and *XIF*^*WT*^) were synthesized and recombined into the pDONR221 Gateway vector and the pB7RWG2 expression vector. All the constructs were re-sequenced to confirm the mutation sites and transformed into *Arabidopsis* plants (WT and *myosin xik xif* mutant background) using flower dip method.

### Protein extraction and immunoblot assay

7 days old homozygous T3 seedlings of *XIK*^*WT*^, *XIK*^*S1234A*^, *XIF*^*WT*^ and *XIF*^*S1256A*^ expressing lines were treated with DMSO or 10 μM NAA for 30 minutes and full seedlings were harvested. Samples were frozen in liquid nitrogen, ground into powder and homogenized in protein extraction buffer (50 mM Tris-HCl, pH=7.5; 150 mM NaCl; 0.15% NP40; 10 mM DTT (1, 4-dithiothreitol); 1 mM PMSF; containing protease inhibitor and phosphatase inhibitor). After 20 minutes of centrifugation (16,000 *g*) at 4 °C, total protein extract was collected and concentration was determined by Bradford Reagent (Bio-Rad). Lambda protein phosphatase treatment (New England BioLabs) was performed following the manufactorer’s protocol. After adding an equal amount of loading buffer (4× Laemmli Sample Buffer, Bio-Rad), an equivalent of 30 μg total protein for each sample was boiled at 65°C for 10 minutes and then separated on a 10% normal or the Phos-tag SDS-PAGE (Phos-tag acrylamide AAL-107, Wako Pure Chemical Industries, #304-93521). Proteins were transferred to a PVDF membrane by wet blotting. The membranes were incubated with primary rat monoclonal anti-5F8 antibody (Chromotek, 1:1000) and secondary bovine anti-rat horseradish peroxidase (HRP)-conjugated (1:5000) antibody. Following detection of the myosin protein, the membranes were stripped and incubated with primary plant Monoclonal Anti-Actin (Sigma, A0480, 1:1000) and secondary anti-mouse HRP (1:500) antibody overnight. HRP activity was detected by the Supersignal Western Detection Reagents (Thermo Scientific) and imaged with a GE Healthcare Amersham 600RGB system.

### Quantification of hypocotyl gravitropic bending angle

To monitor gravitropic responses, plates with 3 days old etiolated seedlings were turned 90° compared to the original gravitropic vector. The plates were scanned 24 hours after gravistimulation and bending was recorded at 1 hour intervals. The hypocotyl bending angle was measured using ImageJ (NIH; http://rsb.info.nih.gov/ij).

### Quantitative analysis of PIN3-GFP polarization in hypocotyl

The *pPIN3::PIN3-GFP* intensity was measured using ImageJ in the hypocotyl as described previously (Rakusová et al., 2019). After gravity stimulation, the ratio between lower and upper side endodermal cells was calculated. For auxin treatment, 3 days old etiolated seedlings were transferred to new plates with 10 μM of NAA, or the equivalent amount of DMSO for 4 hours in darkness after which the inner/outer ratio was calculated for the relevant endodermal cells.

### Hypocotyl basi-petal auxin transport assay

6 days etiolated seedlings were transferred to new ½ MS plates. 3 replicates of 15 seedlings for each mutant or treatment were used. ^3^H-IAA was added into ½ MS medium (1.25% phyto-agar) to a final concentration of 1.5 μM. From this agar solution, a 5 μl ^3^H-IAA droplets was made and once solidified, this droplet was placed on the top of the hypocotyl from which the cotyledons were excised. During the following 6 hours the seedlings were kept in the dark. 10 μM *N*-1-Naphthylphthalamic acid (NPA) in the droplets was used as a control. After 6 hours, the roots and the top of the hypocotyl, which was in contact with the droplet were removed. The remaining hypocotyl samples were homogenized in liquid nitrogen and incubated overnight in Opti-Fluor Scintillation Cocktail (Perkin Elmer). The amount of ^3^H-IAA taken up by the hypocotyl was measured in a scintillation counter (Hidex 300SL) and this represents basipetal polar auxin transport (PAT). Measurements were performed in three technical and three biological replicates.

### Leaf venation assay

7 days old light grown seedlings were used for leaf venation analysis. Cotyledons were cleared in 4% HCl and 20% methanol for 15 minutes at 65℃, followed by a 15 minutes incubation in 7% NaOH and 70% ethanol at room temperature. Next, seedlings were rehydrated by successive incubations in 70%, 50%, 25%, and 10% ethanol for 5 minutes each, followed by incubation in 25% glycerol and 5% ethanol for 2 days at room temperature. Finally, cotyledons were mounted in 50% glycerol and were monitored by differential interference contrast DIC microscopy (Olympus BX53). The cotyledon venation pattern was evaluated by counting the number of loops and defects such as open loops, extra branches and disconnectivity were scored.

### Regeneration of stem after wounding

For the regeneration experiments, young, approximately 10 cm tall *Arabidopsis thaliana* inflorescence stems were tranversally cut at the base, 3 to 4 mm from the rosette. This transversal incission interrupts the longitudinal continuum of vascular cambium and secondary tissues and hence the ongoing transport. Plants were covered with an artificial weight according to Mazur et al. (2016). Axillary buds growing above the rosette were not removed, thereby mainting a source of endogenous auxin. 7 days after wounding, the stem segments were sectioned to 80 μm native sections using an automated vibratome (Leica VT1200 S, Leica Mycrosystems Ltd., Wetzlar, Germany). These sections were stained with a 0.025 % Toluidine Blue O and regeneration was analyzed in stems by observations using a wide field microscope at 10x magnification (Zeiss Axioscope.A1 ZENAxiocam 105).

### Auxin-induced canalization in wounded inflorescence stems

Arabidopsis plants with young, 10 cm tall inflorescence stems were chosen for exogenous auxin application. Stems were wounded by a transversal incision 3-4mm above the rosette to interrupt the vascular cambium and secondary tissues and hence also the polar, basipetal transport of endogenous auxin. We then applied 10 μM IAA (Sigma-Aldrich, cat. no 15148-2G) in a droplet of lanoline paste below the cut. This droplet was replaced every 2 days to ensure the constant presence of auxin. Samples were collected and the manual longitudinal stem sections were obtained under a NIKON MSZ1500 stereomicroscope. Sections were stained in 0,05% Toluidine Blue O and mounted in a 50% glycerol aqueous solution. Images of these sections were obtained by the Olympus BX43 microscope equipped with an Olympus SC30 Camera. n = 20

### Whole-mount *in situ* immunolozalization

For PIN1 localization in root, 3 days old primary roots were treated for 4h with 10 μM of NAA in liquid ½ MS medium. Immunolozalization and quantificaiton of PIN1 lateralization were carried out as described previously (Sauer et al., 2006). The following antibodies were used: anti-PIN1 (1:1000), secondary goat anti-rabbit antibody coupled Cy3 (Sigma-Aldrich, 1:600). PIN1 localization was monitored by LSM 800 microscopy.

### FM4-64 uptake assay

4 days old light grown seedlings were incubated with 10 μM NAA or the equivalent amount of DMSO for 30 minutes in ½ MS liquid medium. Then 2 μM FM4-64 was added to the medium for 15 minutes and roots were mounted and observed using a LSM-800 Zeiss confocal microscope. Signal intensity at PM and cytoplasm (CP) was measured using ImageJ. The ratio between CP and PM was calculated and statistical analysis was performed in GraphPad Prism.

### BFA treatment assay

To monitor PIN1 trafficking in the root, 3 days old light grown seedlings were pretreated with 10 μM NAA or the equivalent amount of DMSO in ½ MS liquid medium for 30 minutes, and then co-treatment with 50 μM BFA for another 60 minutes. Immunolozalization was carried out as described previously (Sauer et al., 2006). The average number of BFA bodies per root was quantified in the different mutant lines.

### Statistical analysis

All statistical analysis was done in GraphPad Prism (version 8.3.0(538)) and significant differences were determined using Anova Tests – Tukey for multiple comparisons. For all graphs the number of * indicate significance according to the following p-values: ns for p > 0.05, * for p ≤ 0.05, ** for p ≤ 0.01, *** for p ≤ 0.001.

## Supplemental Table 1

Table contains analyzed differentially regulated phosphopeptides (FDR ≤0.05) data of IAA, PEO-IAA and cvxIAA treatments.

---

**Figure S1.**
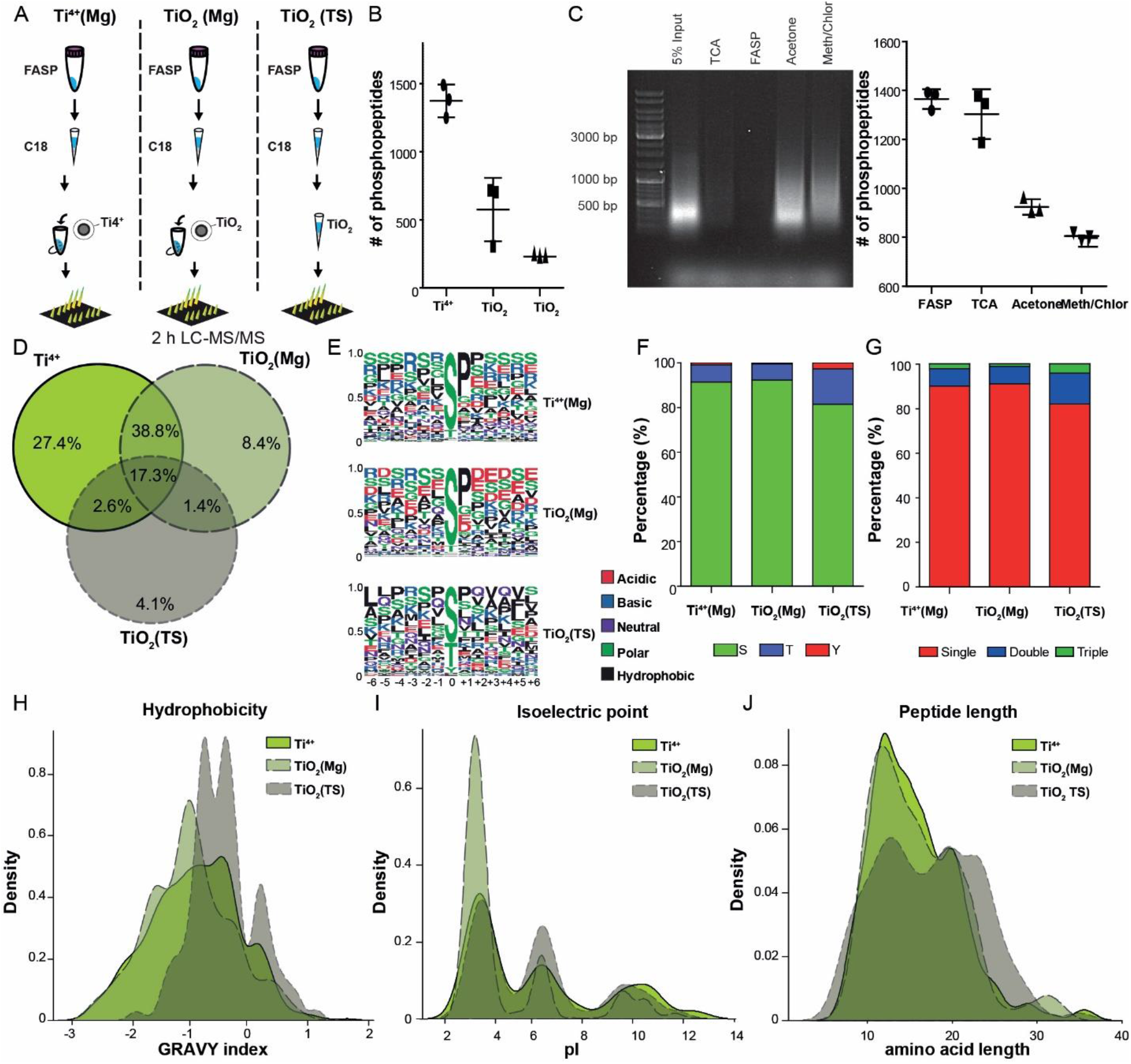
Comparison of different protein precipitation and phospho-peptide enrichments strategies. (A, B) Magnetic IMAC Ti^4+^(Mg), MOAC TiO_2_(Mg) and agarose MOAC TiO_2_(TS) enrichment strategies were compared for phosphopeptide enrichment performance. IMAC outperformed the MOAC methods by identifying ~3 fold more phosphopetides. All strategies were conducted in technical triplicate. (C) In order to analyse nucleic acid interference, to peptides digested samples were compared to 5 % protein lysate input material on a DNA agarose gel. Acetone and methanol/chloroform percipitation techniques retained nucleic acids and performed poorly for the identified number of phosphopeptides. TCA and FASP on the other hand clearly were more optimal precipitation strategies. All samples were handled in technical triplicates and phosphopeptides were enriched by the Ti^4+^-IMAC method. (D) Venn diagram of enriched phosphopeptides showed relatively litlle overlap between the different enrichment methods. Besides having higher numbers of phosphopeptide detection (**Figure S1B**), this graph also indicates that Ti^4+^-IMAC resulted in more uniquely identified phospho-sites. (E) Motif analysis from the specific enriched peptides using the different phosphopeptide enrichment strategies. The color codes of the surrounding amino acids indicate their neutral, acidic, basic, polar or hydrophobic state. The higher presence of acidic amino acids in the TiO_2_(Mg) strategie, indicates a bias of the method towards acidic peptides. (F) The overall nature of phosphorylated amino acids followed a distribution considered normal (Wu et al. 2016) and no discernable differences were observed between the different methods. (G) The charge state of the identified phosphopeptides (namely single, double of triple) did not differ significantly between the different enrichment methods evaluated. (H-J) The biochemical properties of specific peptides confirmed the bias in acidic peptides for TiO_2_(Mg) (lower pI in **Figure S1I**), but the hydrophobility and peptide length of the identified peptides were rather comparable (**Figure S1H-J**).

**Figure S2.**
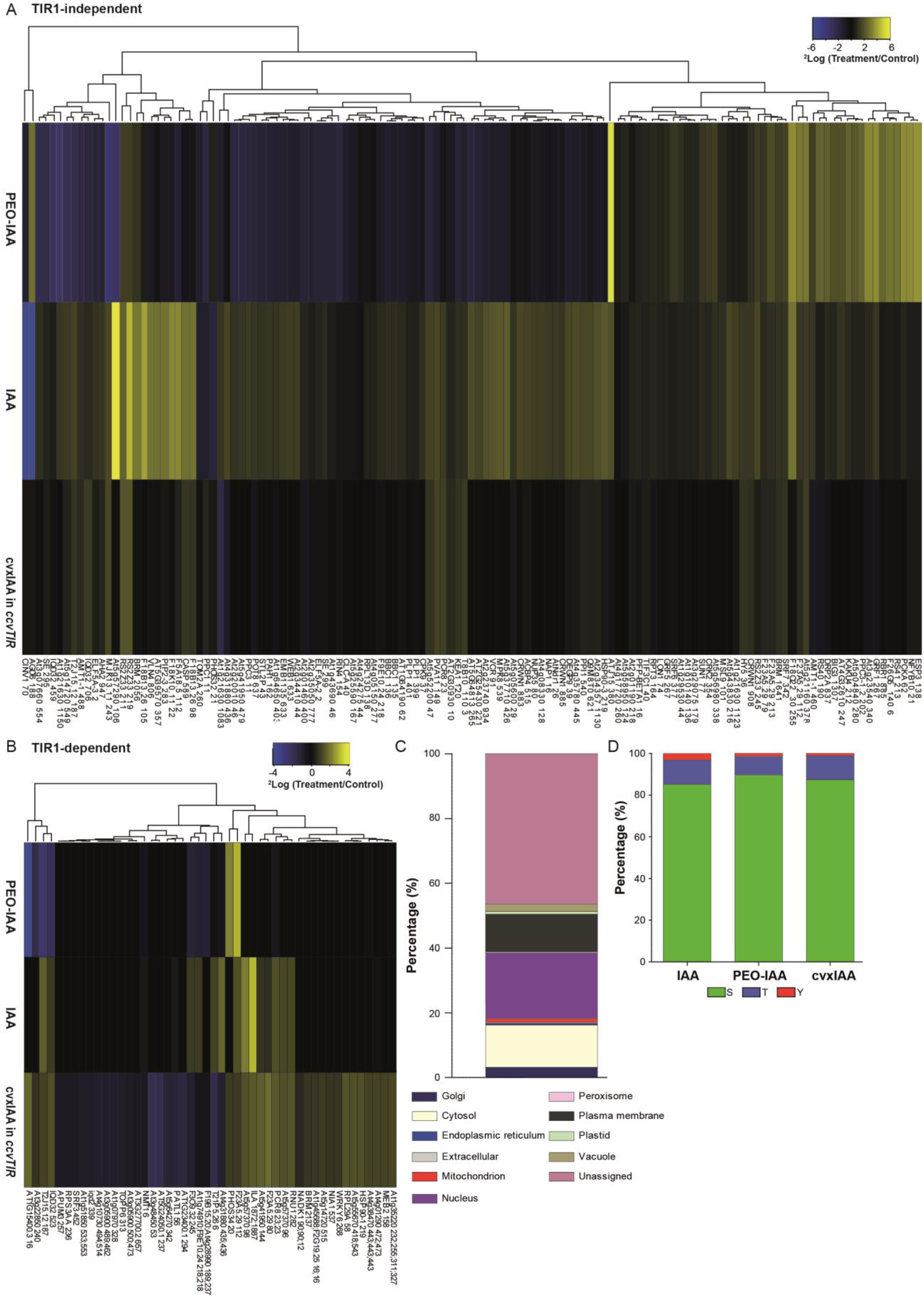
Analysis of phospho-peptide datasets. (A) TIR1-independent and TIR1-dependent rapid auxin response. TIR1-independent, significantly IAA-regulated phospho-peptides, do often show an opposite effect in the PEO-IAA dataset and correspondingly lacked regulation in the cvxIAA dataset (seen by the predominantly black color). (B) The TIR1-dependent, significantly regulated phospho-sites from the cvxIAA treatment did barely overlap with the IAA and PEO-IAA phospho-sites, visualized by the dark colors in these two datasets versus significant up- or downregulation in the cvxIAA sample. (C) The proteins corresponding to differentially regulated phospho-peptides (FDR ≤0.05) from the 100 nM IAA dataset show a predominant localization in the nucleus, cytoplasm and at the plasma membrane. (D) The distribution of phosphorylated amino acids did not significantly differ between IAA, PEO-IAA and cvxIAA treatment and followed the generally expected pattern: Serine(S) ~90%, Threonine(T) ~7 and Tyrosine(Y) ~1% (Wu et al. 2015).

**Figure S3.**
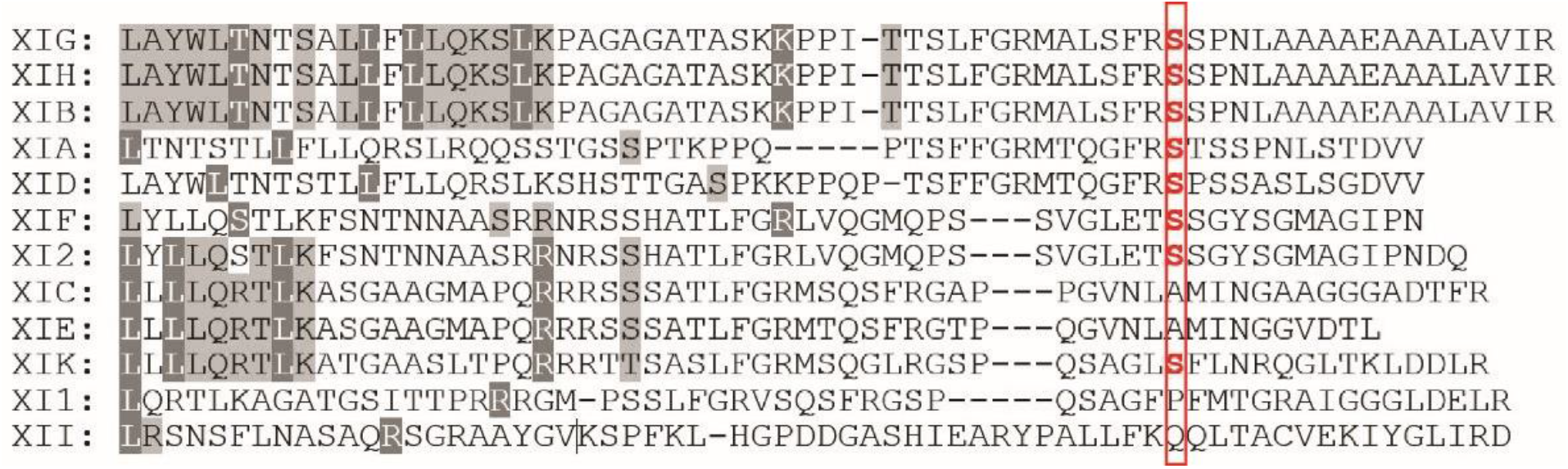
Conserved phosphorylation site in the myosin XI protein family. Protein alignment of the global tail sequence of Myosin XI proteins shows that the auxin-induced phospho-site (S1234 in Myosin XIK) is conserved.

**Figure S4.**
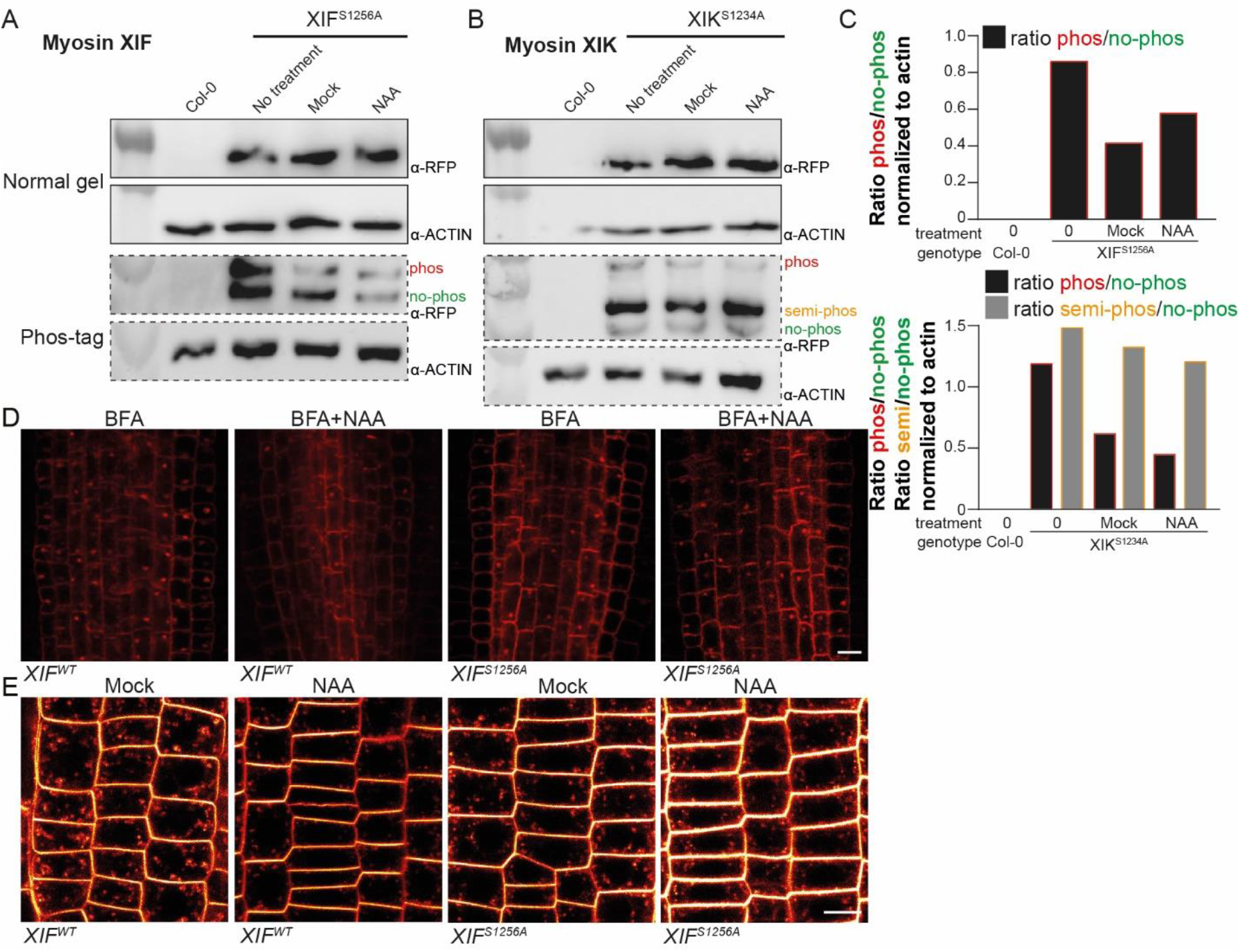
Auxin and trafficking response in phospho-deficient myosin mutants (WT background). (A-C) Phosphorylation of Myosin XI phospho-deficient proteins (resp. *XIF*^*S1256A*^ and *XIK*^*S1234A*^) following auxin treatment visualized by Western and Phos-tag blot assays. 7 days old whole seedlings were treated with 10 μM NAA or mock (DMSO solvent) for 30 minutes. Proteins were extracted of whole seedlings and submitted to Western and Phos-tag blot analysis. The anti-actin loading control within the same sample lane was used to normalize the protein expression level. In these lines, the anti-RFP antibody picks up multiple bands in the Phos-tag gels, for which the intensity was not increased upon auxin treatment as was the case for WT proteins (**Figure 2C-D**). (D) Representative images showing the effect of BFA and the BFA + NAA co-treatment in XIF^WT^ and XIF^S1256A^ mutants in WT background. 3 days old seedlings were pre-treated with 10 μM NAA or DMSO mock for 30 minutes and then co-treated with 50 μM BFA + 10 μM NAA for an additional 60 minutes. Seedlings were immunostained using anti-PIN1 according to Sauer et al. (2006). Scalebar = 10 μm. (E) Representative confocal images of FM4-64 staining in MyosinXIF^WT^ and phospho-deficient XIF^S1256A^ roots (in WT background). Cytoplasmic (CM) and PM staining were observed in 4 days old seedlings treated with 10 μM NAA or DMSO mock for 30 minutes followed by 2 μM FM4-64 staining. Scale bar = 10 μm.

**Figure S5:**
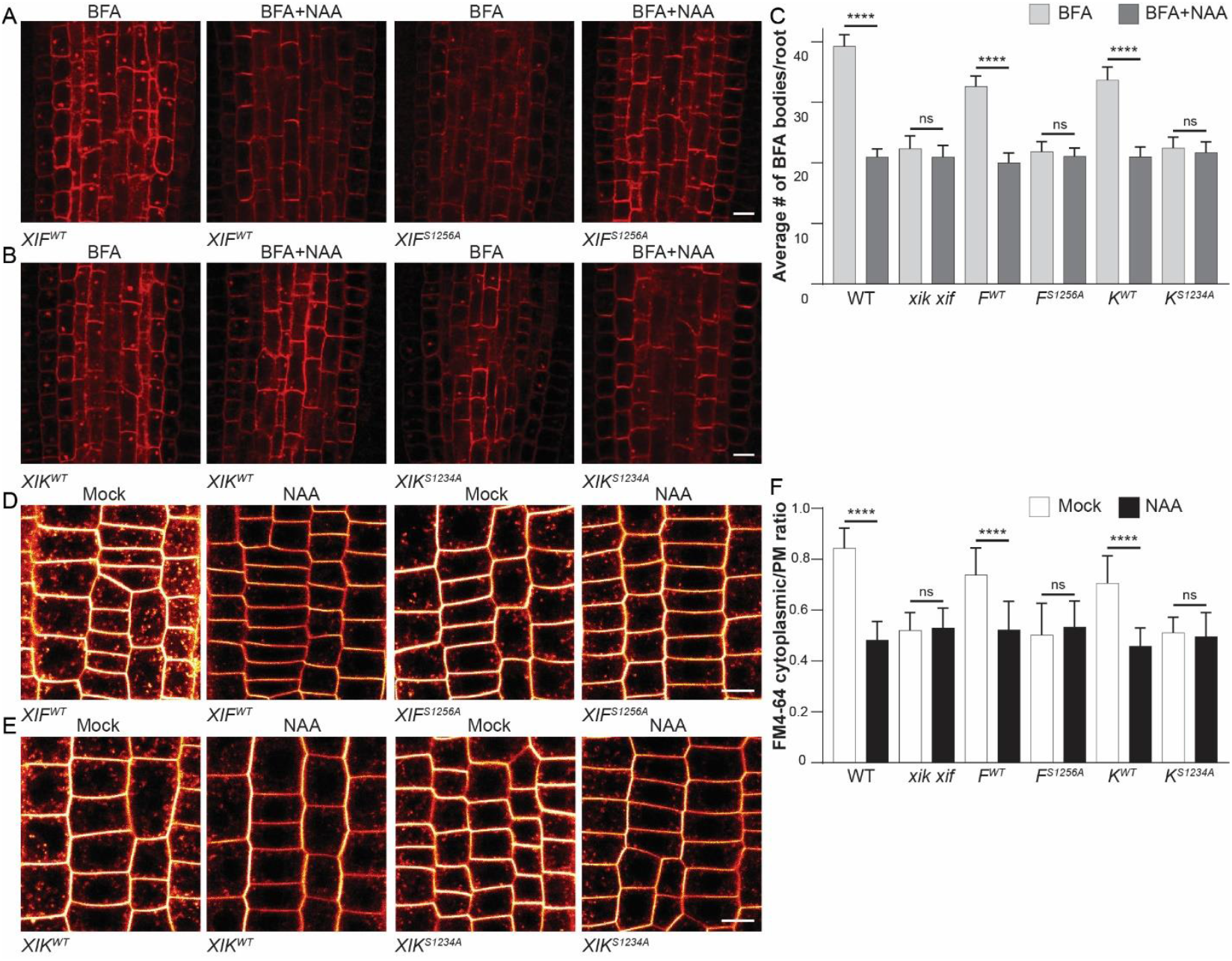
Endomembrante trafficking in phospho-deficient and phospho-mimic mutants (in *myosin xik xif* background) (A) Representative images demonstrating the effect of BFA treatment or the co-treatment BFA + NAA in roots over-expressing XIF^WT^ and XIF^S1256A^ in *myosin xik xif* mutant background. 3 days old seedlings were pre-treated with 10 μM NAA or DMSO mock for 30 minutes and then co-treated with 50 μM BFA and 10 μM NAA for another 60 minutes followed by anti-PIN1 antibody immunostaining according to Sauer et al. (2016). Scale bar = 10 μm. (B) Representative images demonstrating the effect of BFA treatment or the co-treatment BFA + NAA in roots over-expressing XIK^WT^ and XIK^S1234A^ in *myosin xik xif* mutant background. 3 days old seedlings were pre-treated with 10 μM NAA or DMSO mock for 30 minutes and then co-treated with 50 μM BFA and 10 μM NAA for another 60 minutes followed by anti-PIN1 antibody immunostaining according to Sauer et al. (2016). Scale bar = 10 μm. (C) Quantification of BFA bodies in WT, *myosin xik xif* and phospho-deficient XIF^S1256A^ and XIK^S1234A^ roots (**Figure S5A-B**). The average number of BFA bodies was determined per root, including a correction for the total number of cells evaluated per root. n > 20. Bars represent the mean ± SD. **** for p ≤ 0.0001 and ns for p >0.05 determined by Tukey’s multiple comparison test. (D) Representative confocal images of FM4-64 staining in roots over-expressing XIF^WT^ and XIF^S1256A^ (in *myosin xik xif* background). PM staining and internalization of FM4-64 staining was imaged in 4 days old seedlings treated with 10 μM NAA or DMSO mock for 30 minutes. Scale bar = 10 μm. (E) Representative confocal images of FM4-64 staining in roots over-expressing XIK^WT^ and XIK^S1234A^ (in *myosin xik xif* background). PM staining and internalization of FM4-64 staining was imaged in 4 days old seedlings treated with 10 μM NAA or DMSO mock for 30 minutes. Scale bar = 10 μm. (F) Quantification of FM4-64 staining in WT, *myosin xik xif* and all phospho-deficient roots (**Figure S5C-D**). The CM/PM ratio was calculated by dividing the internal staining intensity by the PM-intensity. Bars represent the mean ± SD. Observations were made in more than 50 cells in more than10 different roots. *** for p ≤ 0.001 and **** for p ≤0.0001 determined by Tukey’s multiple comparison test.

**Figure S6.**
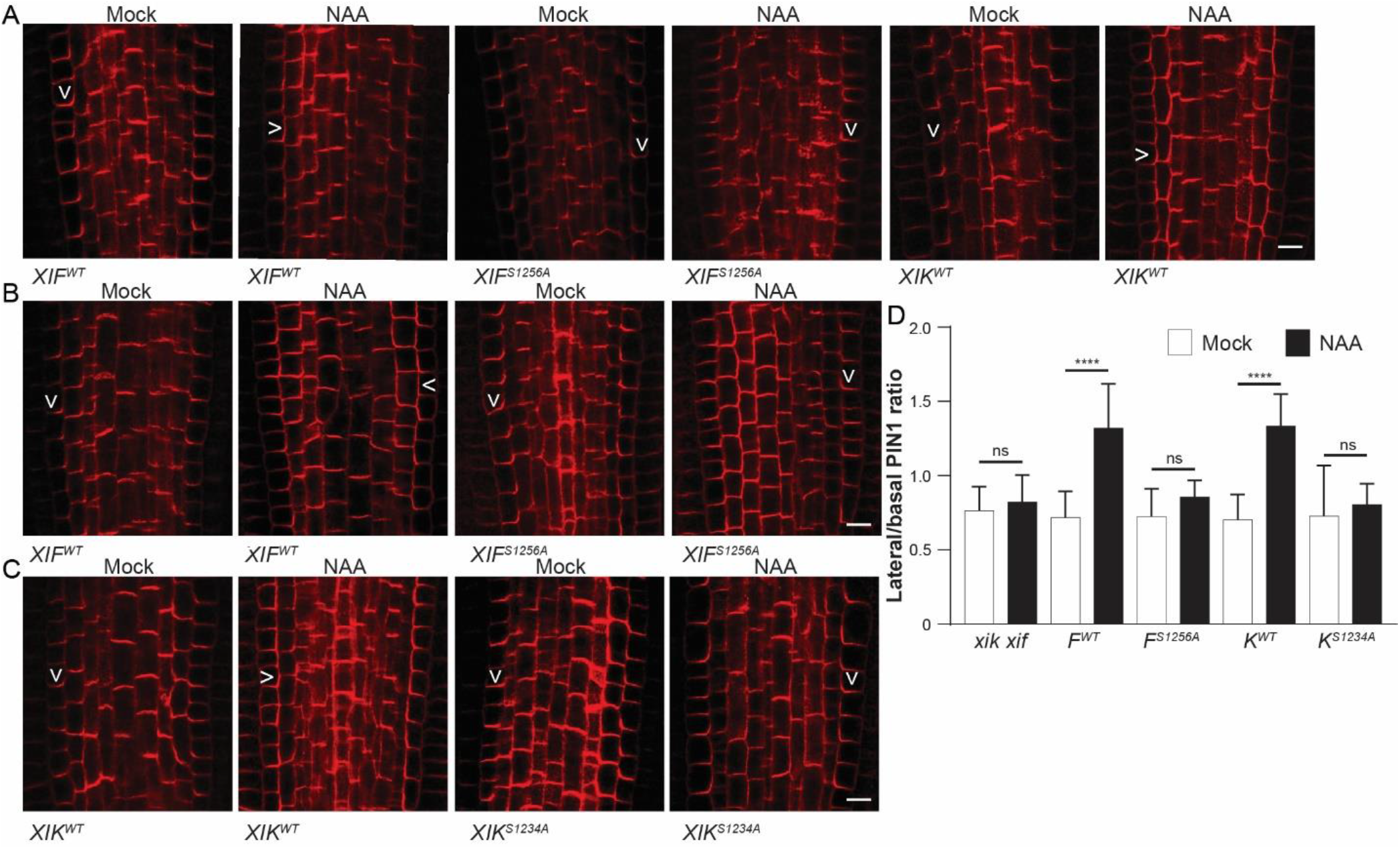
Auxin-induced PIN1 lateralization in WT and phospho-deficient MyosinXI constructs (in WT and in *myosin xik xif* background). (A) PIN1 imunolocalization in WT roots transformed with resp. XIF^WT^, XIF^S1256A^ and XIK^WT^. Arrowheads indicate the predominant PIN1 localization in the endodermal cells. Scale bar = 10 μm. (B) PIN1 imunolocalization in *myosin xik xif* mutant roots transformed with resp. XIF^WT^ and the phospho-deficient Myosin XIF construct (XIF^S1256A^). Arrowheads indicate the predominant PIN1 localization in the endodermal cells. Scale bar = 10 μm. (C) PIN1 imunolocalization in *myosin xik xif* mutant roots transformed with resp. XIK^WT^ and the phospho-deficient Myosin XIK construct (XIK^S1234A^). Arrowheads indicate the predominant PIN1 localization in the endodermal cells. Scale bar = 10 μm. (D) Quantification of auxin-mediated PIN1 lateralization in *myosin xik xif* mutants transformed with the different non-mutated and phospho-deficient MyosinXIF and XIK constructs (**Figure S6B-C**). The ratio was calculated by dividing the PIN1 intensity signal of the lateral side of versus the basal side of the respective endodermal cells. Data represent the mean ± SD. n = 15, more than 50 cells were quantified. **** p ≤0.0001 determined by Tukey’s multiple comparison test.

**Figure S7.**
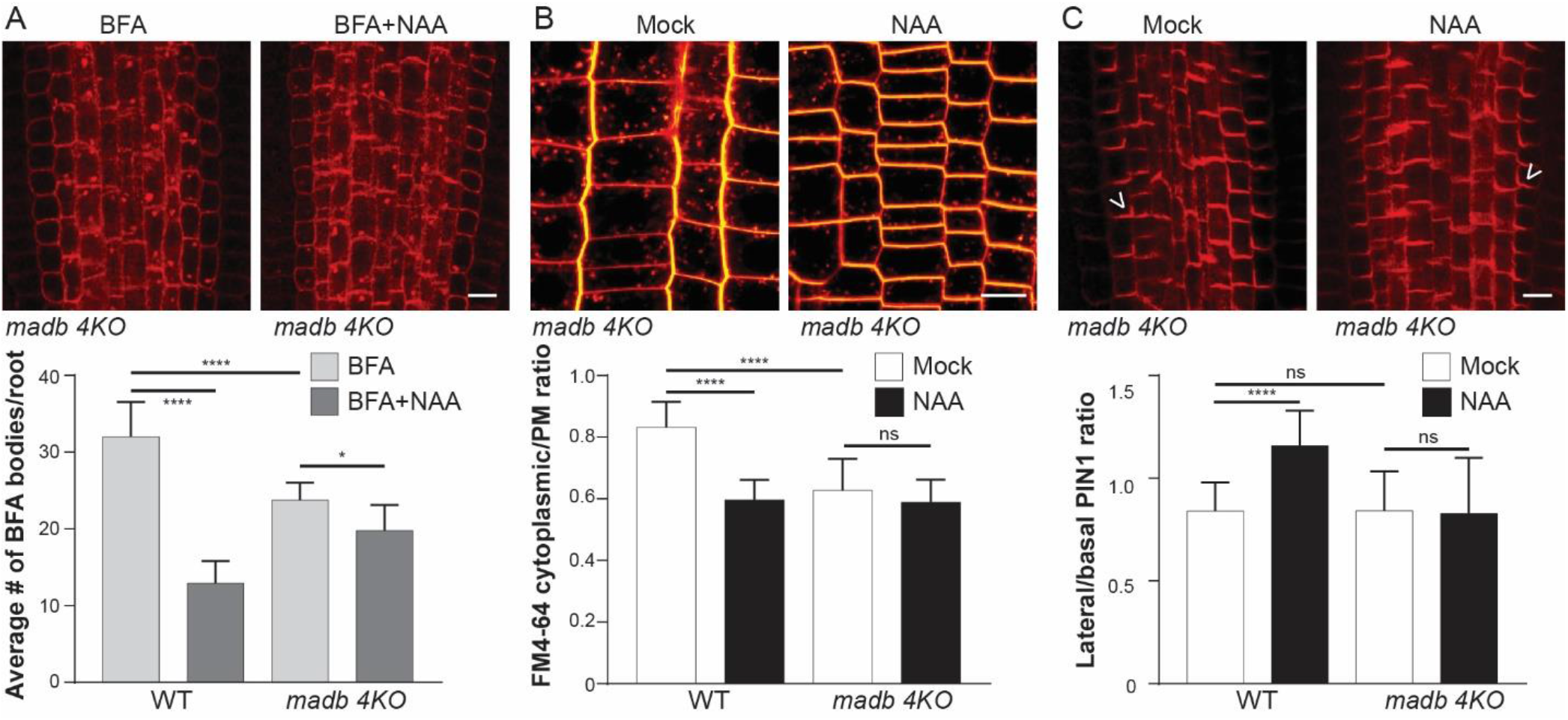
The Madb2 myosin binding protein contributes to PIN trafficking and polarity regulation. (A) Representative images and quantification demonstrating the effect of BFA treatment or the co-treatment BFA + NAA in *madb2 4KO* mutant roots. 3 days old seedlings were pre-treated with 10 μM NAA or DMSO mock for 30 minutes and then co-treated with 50 μM BFA and 10 μM NAA for another 60 minutes followed by anti-PIN1 antibody immunostaining according to Sauer et al. (2016). Scale bar = 10 μm. The average number of BFA bodies was determined per root, including a correction for the total number of cells evaluated per root. n > 20. Average # of BFA bodies per root ± SD is represented. **** for p ≤ 0.0001 and ns for p >0.05 determined by Tukey’s multiple comparison test. (B) Representative confocal images and quantification of FM4-64 staining in *madb2 4KO* roots. PM staining and CM internalization of FM4-64 staining were imaged in 4 days old seedlings treated with 10 μM NAA or DMSO mock for 30 minutes. Scale bar = 10 μm. The CM/PM ratio was calculated by dividing the internal staining intensity by the PM-intensity. Bars represent the mean ± SD. *** for p ≤ 0.001 and **** for p ≤0.0001 determined by Tukey’s multiple comparison test. (C) Representative images and quantification of PIN1 localization in *madb 4ko* mutant roots upon 10 μM NAA-treatment. To visualize the auxin-induced lateralization, the intensity ratio of lateral to basal PIN1 signal was calculated per cell. Whereas 10 μM NAA in WT resulted in lateralization of PIN1 in root endodermal cells, this effect was missing in *madb 4KO* mutant roots. Data represent the mean ± SD. More than 50 cells were evaluated in at least 15 independent roots. **** p ≤0.0001 determined by Tukey’s multiple comparison test. Scale bar = 10 μm

**Figure S8.**
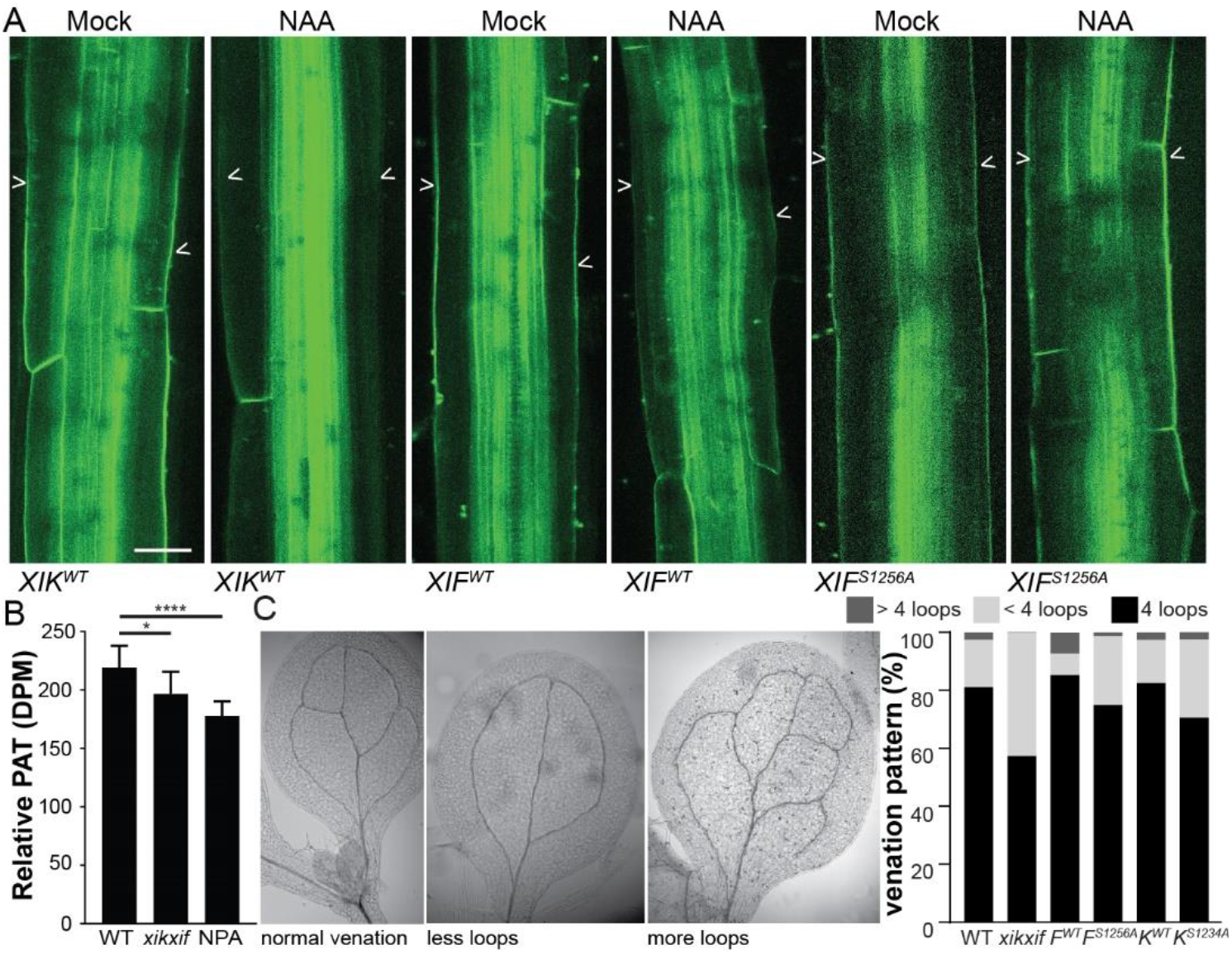
Auxin-induced PIN3 polarization in phospho-deficient myosin mutants in WT background. (A) Representative images showing PIN3-GFP localization hypocotyls expressing the non-mutated Myosin XIF/XIK and the phospho-deficient MyosinXIF^S1256A^ and MyosinXI^KS1234A^ constructs upon DMSO or NAA treatment. Arrowheads indicate PIN3 at outer side of hypocotyl endodermal cells. Scale bar = 25 μm. (B) Polar auxin transport (PAT) in *myosin xik xif* mutant hypocotyls is decreased in comparison to WT, but not as significant as in the NPA-treated hypocotyls. Averages of 3 independent biological repeats existing of 15 hypocotyls ± SD are represented. * p ≤ 0.05 and **** p ≤0.0001 determined by Student t-tests compared to WT.(C) Representative seedlings showing gravity-induced hypocotyl bending of the WT MyosinXIF and XIK constructs compared to their phospho-deficient versions, resp. XIF^S1256A^ and XIK^S1234A^ expressed in *myosin xik xif* mutant background. (C) Representative images of cotyledon vasculature patterns showing the normal venation pattern with 4 loops, less loops and cotyledons with more loops. Scoring of the number of loops was performed in cotyledons of WT, *myosin xik xif* and the different phospho deficient mutants (WT background). n > 50

**Figure S9.**
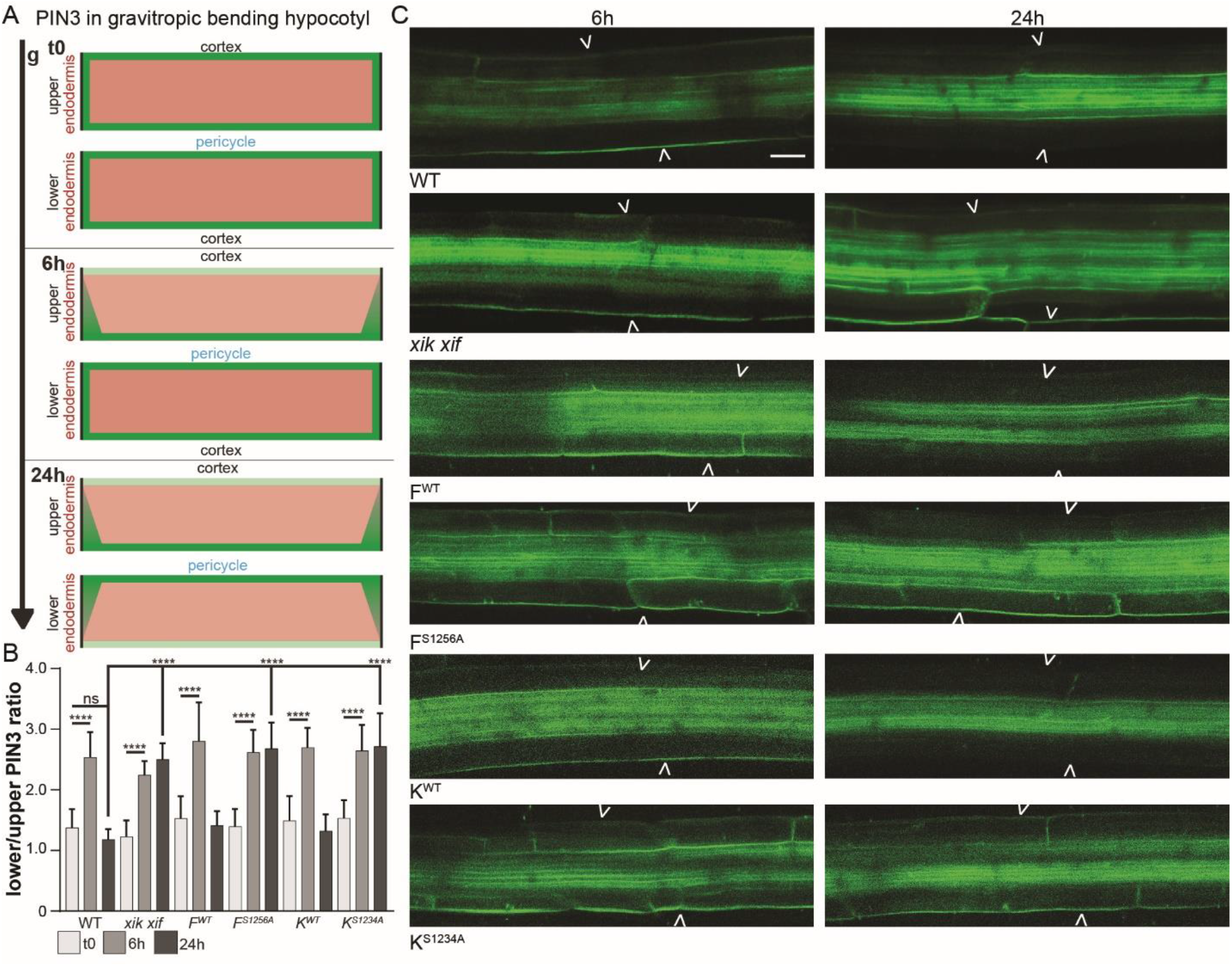
Gravity-induced PIN3 polarization in myosin mutants. (A) Schematic changes of PIN3 in the hypocotyl upon gravity stimulation. (B) Quantification of the bending response in WT, *myosin xik xif* and the lines expressing XIF^WT^/XIK^WT^ and the phospho-deficient mutant versions (XIF^S1256A^ and XIK^S1234A^). PIN3-GFP intensity at the lower and upper side of hypocotyl endodermal cells is represented. The start ratio at t0 is compared to the ratio after 6 and 24h of gravistimulation. At least 20 hypocotyls were evaluated **** p ≤0.0001 determined by Tukey’s multiple comparison test. (C) Representative images of PIN3-GFP localization in gravity-stimulated WT and *myosin xik xif* hypocotyls, 6h and 24h after the gravity switch. In all genotypes the 6h response was normal, with an increased lower/upper endodermal PIN3-GFP ratio. However, in 24h gravity-stimulated *myosin xik xif* hypocotyls and hypocotyls expressing the phospho-deficient XIF^S1256A^ and XIK^S1234A^ versions, no re-establishment of PIN3-GFP balance was observed, indicating that there are problems in the termination of the gravity response, explaining the hyperbending observed in these lines (**Figure 6B**). Arrowheads indicate the outer side of hypocotyl endodermal cells. Scale bar = 20 μm.

## References

Abel, S., and Theologis, A. (1996). Early genes and auxin action. Plant Physiol. 111, 9

Abas, L., Benjamins, R., Malenica, N., Paciorek, T., Wišniewska, J., Moulinier–Anzola, J. C., Sieberer, T., Friml, J. and Luschnig, C. (2006). Intracellular trafficking and proteolysis of the Arabidopsis auxin-efflux facilitator PIN2 are involved in root gravitropism. Nat. Cell Biol. 8, 249–256

Abu-Abied, M., Belausov, E., Hagay, S., Peremyslov, V., Dolja, V. and Sadot, E. (2018). Myosin XI-K is involved in root organogenesis, polar auxin transport, and cell division. J. Exp. Bot. 69, 2869–2881

Adamowski, M., and Friml, J. (2015). PIN-dependent auxin transport: action, regulation and evolution. Plant Cell 27, 20–32

Avisar D., Prokhnevsky, A. I., Makarova, K. S., Koonin, E. V. and Dolja, V. V. (2008) Myosin XI-K is required for rapid trafficking of golgi stacks, peroxisomes, and mithochondria in leaf cells of *Nicotiana benthamiana*. Plant Physiol. 146: 1098–108

Avisar, D., Abu-Abied, M., Belausov, E., Sadot, E., Hawes, C. and Sparkes, I.A. (2009). A comparative study of the involvement of 17 *Arabidopsis* myosin family members on the motility of Golgi and other organelles. Plant Physiol. 150, 700–709

Barbosa, I.C.R., Hammes, U.Z. and Schwechheimer, C. (2018). Activation and polarity control of PIN-FORMED auxin transporters by phosphorylation. Trends in Plant Sci. 23, 523–538

Bargmann, OR B., Vanneste, S., Krouk, G., Nawy,T., Efroni, I., Shani E., Choe, G., Friml, J., Bergmann C D., Estelle, M. and Birnbaum, D K. (2013) A map of cell type-specific auxin response. Mol. Syst. Biol. 9: 688

Benková, E., Michniewicz, M., Sauer, M., Teichmann, T., Seifertová, D., Jürgens, G. and Friml, J. (2003). Local, efflux-dependent auxin gradients as a common module for plant organ formation. Cell 115, 591–602

Bennett, T., Hines, G. and Leyser, O. (2014). Canalization: what the flux? Trends in Gen. 30, 41–48

Bhatia, N., Bozorg, B., Larsson, A., Ohno, C., Jönsson, H. and Heisler, M. G. (2016). Auxin acts through MONOPTEROS to regulate plant cell polarity and pattern phyllotaxis. Curr. Biol. 26, 3202–3208

Cao, M., Chen, R., Li, P., Yu, Y., Zheng, R., Ge, D., Zheng, W., Wang, X., Gu, Y., Gelova, Z., Friml, J., Zhang, H., Liu, R., He, J. and Xu, T. (2019). TMK1-mediated auxin signalling regulates differential growth of the apical hook. Nature 568, 240–243

Dhonukshe, P., Aniento, F., Hwang, I., Robinson, D. G., Mravec, J., Stierhof, Y. D. and Friml, J. (2007). Clathrin-mediated constitutive endocytosis of PIN auxin efflux carriers in *Arabidopsis*. Curr. Biol. 17, 520–527

Dindas, J., Scherzer, S., Roelfsema, M.R.G., von Meyer, K., Muller, H.M., Al-Rascheid, K.A.S., Palme, K., Dietrich, P., Becker, D., Bennett, M.J. and Hedrich, R. (2018). AUX1-mediated root hair auxin influx governs SCF^TIR1/AFB^-type Ca^2+^ signaling. Nat. Comm. 9, 1174

Fendrych, M., Akhmanova, M., Merrin, J., Glanc, M., Hagihara, S., Takahashi, K. and Friml, J. (2018). Rapid and reversible root growth inhibition by TIR1 auxin signaling. Nat. Plants 4, 453–459

Gallei, M., Luschnig, C. and Friml, J. (2020). Auxin signaling in growth: Schrödinger’s cat out of the bag. Curr. Opin. Plant Biol. 53, 43–49

Geldner, N., Friml, J., Stierhof, Y. D., Jürgens, G. and Palme, K. (2001). Auxin transport inhibitors block PIN1 cycling and vesicle trafficking. Nature 413, 425

Glanc, M., Fendrych, M. and Friml, J. (2018). Mechanistic framework for cell-intrinsic re-establishment of PIN2 polarity after cell division. Nat. Plants 4, 1082–1088

Govindaraju, P., Verna, C., Zhu, T. and Scarpella, E. (2020). Vein patterning by tissue-specific auxin transport. Development 147, dev187666

Hajný, J., Prát, T., Rydza, N., Rodriguez, L., Tan, S., Verstraeten, I., Domjan, D., Mazur, E., Smakowska-Luzan, E., Smet, W., Mor, E., Nolf, J., Yang, B.J., Grunewald, W., Molnár, G., Belkhadir, Y., De Rybel, B. and Friml, J. (2020). Receptor kinase module targets PIN-dependent auxin transport during canalization. Science 370, 550–557

Han, H., Rakusova, H., Verstraeten, I., Zhang, Y. and Friml, J. (2020). SCF^TIR1/AFB^ auxin signaling for bending termination during shoot gravitropism. Plant Physiol. 183, 37–40

Haraguchi, T., Ito, K., Duan, Z., Rula, S., Takahashi, K., Shibuya, Y., Hagino, N., Miyatake, Y., Nakano, A., Tominaga, M. (2018). Functional diversity of class XI Myosins in *Arabidopsis thaliana*. Plant Cell Physiol. 29: 2268–2277

Hayashi, K.I., Tan, X., Zheng, N., Hatate, T., Kimura, Y., Kepinski, S. and Nozaki, H. (2008). Small-molecule agonists and antagonists of F-box protein-substrate interactions in auxin perception and signaling. Proc. Natl. Acad. Sci. USA 105, 5632–5637

Heisler, M. G., Ohno, C., Das, P., Sieber, P., Reddy, G. V., Long, J. A. and Meyerowitz, E. M. (2005). Patterns of auxin transport and gene expression during primordium development revealed by live imaging of the Arabidopsis inflorescence meristem. Curr. Biol. 15, 1899–1911

Hooper, C. M., Castleden, I. R., Tanz, S. K., Aryamanesh, N. and Millar, A. H. (2017). SUBA4: the interactive data analysis centre for Arabidopsis subcellular protein locations. Nucleic Acids Res. 45, D1064–D1074

Humphrey, S. J., Azimifar, S. B. and Mann, M. (2015). High-throughput phosphoproteomics reveals *in vivo* insulin signaling dynamics. Nat. Biotechnol. 33, 990–995.

Jásik, J., Bokor, B., Stuchlík, S., Mičieta, K., Mičieta, K., Schmelzer, E. (2016). Effects of auxin on PIN-FORMED2 (PIN2) dynamics are not mediated by inhibiting PIN2 endocytosis. Plant Physiol. 172, 1019–1031

Jelínková, A., Malínská, K., Simon, S., Kleine-Vehn, J., Parezová, M., Pejchar, P., Kubes, M., Martinec, J., Friml, J., Zazímalová, E. and Petrásek J. (2010). Probing plant membranes with FM dyes: tracking, dragging or blocking? Plant J. 61, 883–892

Kieffer, M., Neve, J. and Kepinski, S. (2010). Defining auxin response contexts in plant development. Curr. Opin. Plant Biol. 13, 12–20

Kleine-Vehn, J., Leitner, J., Zwiewka, M., Sauer, M., Abas, L., Luschnig, C. and Friml, J. (2008a). Differential degradation of PIN2 auxin efflux carrier by retromer-dependent vacuolar targeting. Proc. Natl. Acad. Sci. USA 105, 17812–17817

Kleine-Vehn, J., Langowski, L., Wisniewska, J., Dhonukshe, P., Brewer, P.B., Friml J. (2008b). Cellular and molecular requirements for polar targeting and transcytosis in plants. Mol. Plant 1, 1056–1066

Koonin, E. V. and Dolja, V. V. (2017). Myosin-driven transport network in plants. Proc. Natl. Acad. Sci. USA 114, E1385–E1394

Kubeš, M. and Napier, R. (2019). Non-canonical auxin signalling: fast and curious. J. Exp. Bot. 70, 2609

Kurth, E. G., Peremyslov, V. V., Turner, H. L., Makarova, K. S., Iranzo, J., Mekhedov, S. L., Lavy, M. and Estelle, M. (2016) Mechanisms of auxin signaling. Development 143, 3226–3229

McClure, B. A., Hagen, G., Brown, C. S., Gee, M. A. and Guilfoyle, T. J. (1989). Transcription, organization, and sequence of an auxin-regulated gene cluster in soybean. Plant Cell, 1, 229–239

Lewis, D. R., Olex, A. L., Lundy, S. R., Turkett, W. H., Fetrow, J. S. and Muday, G. K. (2013). A kinetic analysis of the auxin transcriptome reveals cell wall remodeling proteins that modulate lateral root development in Arabidopsis. Plant Cell 25, 3329–3346

Li, L., Verstraeten, I., Roosjen, M., Takahashi, K., Rodriguez, L., Merrin, J., Chen, J., Shabala, L., Smet, W., Ren, H., Vanneste, S., Shabala, S., De Rybel, B., Weijers, D., Kinoshita, T., Gray, W.M. and Friml, J. (2021). Antagonistic cell surface and intracellular auxin signalling regulate plasma membrane H+-fluxes for root growth. Research Square DOI: 10.21203/rs.3.rs-266395/v1

Lin, W., Tang, W., Takahashi, K., Ren, H., Pan, S., Zheng, H., Gray, W.M., Xu, T., Kinoshita, T. and Zhenbiao, Y. (2021). TMK-based cell surface auxin signaling activates cell wall acidification in Arabidopsis. Research Square DOI: 10.21203/rs.3.rs-203621/v1

Mazur, E., Benková, E. and Friml, J. (2016). Vascular cambium regeneration and vessel formation in wounded inflorescence stems of Arabidopsis. Sci. Rep. 6, 33754

Mazur, E., Gallei, M., Adamowski, M., Han, H., Robert, H. S. and Friml, J. (2020a). Clathrin-mediated trafficking and PIN trafficking are required for auxin canalization and vascular tissue formation in Arabidopsis. Plant Sci. 293, 110414

Mazur, E., Kulik, I., Hajný, J. and Friml, J. (2020b). Auxin canalization and vascular tissue formation by TIR1/AFB‐mediated auxin signaling in Arabidopsis. New Phytol. 226, 1375–1383

Michniewicz, M., Zago, M. K., Abas, L., Weijers, D., Schweighofer, A., Meskiene, I., Heisler, M. G., Ohno, C., Zhang, J., Huang, F., Schwab, R., Weigel, D., Meyerowitz, E. M., Luschnig, C., Offringa, R., Friml, J. (2007). Antagonistic regulation of PIN phosphorylation by PP2A and PINOID directs auxin flux. Cell 130, 1044–1056

Morffy, N., and Strader, L. C. (2020). Old Town Roads: routes of auxin biosynthesis across kingdoms. Curr. Opin. Plant Biol. 55, 21–27

Mutte, K. S., Kato, H., Rothfels,. C., Melkonian, M., Wong, G. K. and Weijers, D. (2018). Origin and evolution of the nuclear auxin response system. Elife 7: e33399

Narasimhan, M., Johnson, A., Prizak, R., Kaufmann, W.A., Tan, S., Casillas-Pérez, B. and Friml, J. (2020). Elife 9: e52067

Nikonorova, N., Van den Broeck, L., Zhu, S., Van De Cotte, B., Dubois, M., Gevaert, K., Inze, D. and De Smet, I. (2018). Early mannitol-triggered changes in the Arabidopsis leaf (phospho) proteome reveal growth regulators. J. Exp. Bot. 69, 4591–4607

Okamoto, K., Ueda, H., Shimada, T., Tamura, K., Kato, T., Tasaka, M., Morita, M. T. and Hara-Nishimura, I. (2015). Regulation of organ straightening and plant posture by an actin–myosin XI cytoskeleton. Nat. Plants 1, 15031

Ojangu, E. L., Ilau, B., Tanner, K., Talts, K., Ihoma, E., Dolja, V. V., Paves, H. and Truve, E. (2018). Class XI myosins contribute to auxin response and senescence-induced cell death in *Arabidopsis*. Front. Plant Sci. 9, 1570

Oochi, A., Hajny, J., Fukui, K., Nakao, Y., Gallei, M., Quareshy, M., Takahashi, K., Kinoshita, T., Harborough, S.R., Kepinski, S., Kasahara, H., Napier, R., Friml, J. and Hayashi, K. (2019). Pinstatic acid promotes auxin transport by inhibiting PIN internalization. Plant Physiol. 180, 1152–1165

Paciorek, T., Zažímalová, E., Ruthardt, N., Petrášek, J., Stierhof, Y. D., Kleine-Vehn, J., Morris, A. D., Emans, N., Jürgens, G., Geldner, N. and Friml, J. (2005). Auxin inhibits endocytosis and promotes its own efflux from cells. Nature 435, 1251

Peremyslov, V. V., Prokhnevsky, A. I., Avisar, D. and Dolja, V. V. (2008). Two class XI myosins function in organelle trafficking and root hair development in Arabidopsis. Plant Physiol. 146, 1109–1116

Peremyslov, V.V., Klocko, A. L., Fowler, J. E. and Dolja, V. V. (2012). Arabidopsis Myosin XI-K localizes to the motile endomembrane vesicles associated with F-actin. Front. Plant Sci. 3: 184

Peremyslov, V. V., Morgun, E. A., Kurth, E. G., Makarova, K. S., Koonin, E. V. and Dolja, V. V. (2013). Identification of myosin XI receptors in Arabidopsis defines a distinct class of transport vesicles. Plant Cell 25, 3022–3038

Petrášek, J., Mravec, J., Bouchard, R., Blakeslee, J. J., Abas, M., Seifertová, D., Wisniewska, J., Tadele, Z., Kubes, M., Covanova, M., Dhonukshe, P., Skupa, P., Benkova, E., Perry, L., Krecek, P., Lee, O.R., Fink, G.R., Geisler, M., Murphy, A.S., Luschnig, C., Zazimalova, E.and Friml, J. (2006). PIN proteins perform a rate-limiting function in cellular auxin efflux. Science 312, 914–918

Prát, T., Hajný, J., Grunewald, W., Vasileva, M., Molnár, G., Tejos, R., Schmid, M, Sauer, M. and Friml, J. (2018). WRKY23 is a component of the transcriptional network mediating auxin feedback on PIN polarity. PLoS Genet. 14, e1007177

Prokhnevsky, A. I., Peremyslov, V. V. and Dolja, V. V. (2008). Overlapping functions of the four class XI myosins in *Arabidopsis* growth, root hair elongation, and organelle motility. Proc. Natl. Acad. Sci. USA 105, 19744–19749

Rakusová, H., Abbas, M., Han, H., Song, S., Robert, H. S. and Friml, J. (2016). Termination of shoot gravitropic responses by auxin feedback on PIN3 polarity. Curr. Biol. 26, 3026–3032

Rakusová, H., Gallego‐Bartolomé, J., Vanstraelen, M., Robert, H. S., Alabadí, D., Blázquez, M. A., Benkova, E. and Friml, J. (2011). Polarization of PIN3‐dependent auxin transport for hypocotyl gravitropic response in *Arabidopsis thaliana*. Plant J. 67, 817–826

Rakusová, H., Han, H., Valošek, P. and Friml, J. (2019). Genetic screen for factors mediating PIN polarization in gravistimulated *Arabidopsis thaliana* hypocotyls. Plant J. 98, 1048–1059

Reddy, A. S. and Day, I. S. (2001). Analysis of the myosins encoded in the recently completed *Arabidopsis thaliana* genome sequence. Genome Biol. 2, research0024

Robert, H. S., Grones, P., Stepanova, A. N., Robles, L. M., Lokerse, A. S., Alonso, J. M., Weijers, D. and Friml, J. (2013). Local auxin sources orient the apical-basal axis in Arabidopsis embryos. Curr. Biol. 23, 2506–2512

Robert, S., Kleine-Vehn, J., Barbez, E., Sauer, M., Paciorek, T., Baster, P., Vanneste, S., Zhang, J., Simon, S., Covanova, M., Hayashi, K., Dhonukshe, P., Yang, Z., Bednarek, S.Y., Jones, A.M., Luschnig, C., Aniento, F. and Friml, J. (2010). ABP1 mediates auxin inhibition of clathrin-dependent endocytosis in Arabidopsis. Cell 143, 111–121

Robert, H. S., Park, C., Gutièrrez, C. L., Wójcikowska, B., Pěnčík, A., Novák, O., Chen, J., Grunewald, W., Dresselhaus, T., Friml, J. and Laux, T. (2018). Maternal auxin supply contributes to early embryo patterning in Arabidopsis. Nat. Plants, 4, 548–553

Ryan, J. M. and Nebenführ, A. (2017). Update on myosin motors: molecular mechanisms and physiological functions. Plant Physiol. 176, 119–127

Sachs, T. (1981). The control of the patterned differentiation of vascular tissues. Adv. Bot. Res. 9, 151–162

Salanenka, Y., Verstraeten, I., Löfke, C., Tabata, K., Naramoto, S., Glanc, M. and Friml, J. (2018). Gibberellin DELLA signaling targets the retromer complex to redirect protein trafficking to the plasma membrane. Proc. Natl. Acad. Sci. USA, 115, 3716–3721

Salehin, M., Bagchi, R. and Estelle, M. (2015). SCF^TIR1/AFB^-based auxin perception: mechanism and role in plant growth and development. Plant Cell 27, 9–19

Sauer, M., Balla, J., Luschnig, C., Wiśniewska, J., Reinöhl, V., Friml, J. and Benková, E. (2006). Canalization of auxin flow by Aux/IAA-ARF-dependent feedback regulation of PIN polarity. Genes Dev. 20, 2902–2911

Scarpella, E., Marcos, D., Friml, J. and Berleth, T. (2006). Control of leaf vascular patterning by polar auxin transport. Genes Dev. 20, 1015–1027

Stecker, K.E., Minkoff, B.B. and Sussman, M. R. (2014). Phosphoproteomic analyses reveal early signaling events in the osmotic stress response. Plant Physiol. 165, 1171–1187

Steinman, M.Q. and Trainor, B.C. (2010). Rapid Effects of Steroid Hormones on Animal Behavior. Nature Education Knowledge 3, 1

Talts, K., Ilau, B., Ojangu, E. L., Tanner, K., Peremyslov, V. V., Dolja, V. V., Truve, E. and Paves, H. (2016). Arabidopsis myosins XI1, XI2, and XIK are crucial for gravity-induced bending of inflorescence stems. Front. Plant Sci. 7, 1932

Tan, S., Luschnig, C., Friml, J. (2021) Pho-view of Auxin: Reversible protein phosphorylation in auxin biosynthesis, transport and signaling. Mol. Plant 14, 151–165

Tan, S., Zhang, X., Kong, W., Yang, X. L., Molnár, G., Vondráková, Z., Filepova, R., Petrasek, J., Friml, J. and Xue, H.W. (2020). The lipid code-dependent phosphoswitch PDK1–D6PK activates PIN-mediated auxin efflux in Arabidopsis. Nat. Plants 6, 556–569

Uchida, N., Takahashi, K., Iwasaki, R., Yamada, R., Yoshimura, M., Endo, T.A., Kimura, S., Zhang, H., Nomoto, M., Tada, Y., Kinoshita, T., Itami, K., Hagihara, S. and Torii, K.U. (2018). Chemical hijacking of auxin signaling with an engineered auxin-TIR1 pair. Nat. Chem. Biol. 14, 299–305

Vanneste, S. and Friml, J. (2009). Auxin: a trigger for change in plant development. Cell 136, 1005–1016

Vu, L. D., Stes, E., Van Bel, M., Nelissen, H., Maddelein, D., Inzé, D. and De Smet, I. (2016). Up-to-date workflow for plant (phospho) proteomics identifies differential drought-responsive phosphorylation events in maize leaves. J. of Proteome Res. 15, 4304–4317

Wabnik, K., Govaerts, W., Friml, J. and Kleine-Vehn, J. (2011). Feedback models for polarized auxin transport: an emerging trend. Mol. Syst. Biol. 7, 2352–2359

Wabnik, K., Kleine‐Vehn, J., Balla, J., Sauer, M., Naramoto, S., Reinöhl, V., Merks, R.M.H., Govaerts, W. and Friml, J. (2010). Emergence of tissue polarization from synergy of intracellular and extracellular auxin signaling. Mol. Syst. Biol. 6, 447

Weijers, D. and Wagner, D. (2016). Transcriptional responses to the auxin hormone. Ann. Rev. Plant Biol. 67, 539–574

Wendrich, J. R., Boeren, S., Möller, B. K., Weijers, D. and De Rybel, B. (2017). In vivo identification of plant protein complexes using IP-MS/MS. Methods Mol. Biol. 1497, 147–158

Wiśniewska, J., Xu, J., Seifertová, D., Brewer, P. B., Růžička, K., Blilou, I., Rouquie, D., Benkova, E., Scheres, B. and Friml, J. (2006). Polar PIN localization directs auxin flow in plants. Science 312, 883–883

Wu, L., Hu, X., Wang, S., Tian, L., Pang, Y., Han, Z., Wu, L. and Chen, Y. (2015). Quantitative analysis of changes in the phosphoproteome of maize induced by the plant hormone salicylic acid. Sci. Rep. 5, 1–16

Xiao, Y. and Offringa, R. (2020). PDK1 regulates auxin transport and Arabidopsis vascular development through AGC1 kinase PAX. Nat. Plants 6, 544–555

Xu, T., Dai, N., Chen, J., Nagawa, S., Cao, M., Li, H., Zhou, Z., Chen, Z., De Rycke, R. Rakusova, H., Wang, W., Jones, A.M., Friml, J., Patterson, S.E., Bleecker, A.B. and Yang, Z. (2014). Cell surface ABP1-TMK auxin-sensing complex activates ROP GTPase signaling. Science 343, 1025–1028

Žádníková, P., Petrášek, J., Marhavý, P., Raz, V., Vandenbussche, F., Ding, Z., Schwarzerová, K., Morita, M. T., Tasaka, M., Hejátko, J., Van Der Straeten, D., Friml, J. and Benková, E. (2010). Role of PIN-mediated auxin efflux in apical hook development of *Arabidopsis thaliana*. Development 137, 607–617

Zhang, H., Zhou, H., Berke, L., Heck, A. J., Mohammed, S., Scheres, B. and Menke, F. L. (2013). Quantitative phosphoproteomics after auxin-stimulated lateral root induction identifies an SNX1 protein phosphorylation site required for growth. Mol. Cell Proteomics 12, 1158–1169

Zhang, J., Mazur, E., Balla, J., Gallei, M., Kalousek, P., Medveďová, Z., Li, Y., Wang, Y., Prat, T., Vasileva, M., Reinohl, V., Prochazka, S., Halouzka, R., Tarkowski, P., Luschnig, C., Brewer, P.B. and Friml, J. (2020). Strigolactones inhibit auxin feedback on PIN-dependent auxin transport canalization. Nat. Commun. 11, 1–10

Zhang, J., Nodzyński, T., Pěnčík, A., Rolčík, J. and Friml, J. (2010). PIN phosphorylation is sufficient to mediate PIN polarity and direct auxin transport. Proc. Natl. Acad. Sci. USA 107, 918–922

